# Distance-dependent inhibition of translation initiation by downstream out-of-frame AUGs reveals that ribosome scans in a Brownian ratchet process

**DOI:** 10.1101/2021.11.01.466764

**Authors:** Ke Li, Jinhui Kong, Shuo Zhang, Tong Zhao, Wenfeng Qian

**Affiliations:** State Key Laboratory of Plant Genomics, Institute of Genetics and Developmental Biology, Innovation Academy for Seed Design, Chinese Academy of Sciences, Beijing 100101, China; University of Chinese Academy of Sciences, Beijing 100049, China; Institute of Microbiology, Chinese Academy of Sciences, Beijing 100101, China

**Keywords:** Translation initiation, leaky scanning, downstream AUG, strictly unidirectional scanning model, Brownian ratchet scanning model, preinitiation complex, protein homeostasis

## Abstract

While eukaryotic ribosomes are widely presumed to scan mRNA for the AUG codon to initiate translation in a strictly 5′–3′ movement (strictly unidirectional scanning model), other evidence has suggested that the ribosome uses small-amplitude 5′–3′ and 3′–5′ oscillations with a net 5′–3′ movement to recognize the AUG codon (Brownian ratchet scanning model). Here, we generated 13,437 yeast variants, each with an ATG triplet placed downstream (dATGs) of the annotated ATG (aATG) codon of green fluorescent protein. We found that out-of-frame dATGs could inhibit translation at the aATG, but with diminishing strength over increasing distance between aATG and dATG, undetectable beyond ∼17 nt. Computational simulations revealed that each triplet is scanned back and forth approximately ∼10 times until an AUG codon is recognized. Collectively, our findings uncover the basic process by which eukaryotic ribosomes scan for initiation codons, and how this process could shape eukaryotic genome evolution and influence cancer development.

## Introduction

To synthesize functional proteins and maintain protein homeostasis, the genetic information carried by messenger RNAs (mRNAs) must be faithfully transmitted to proteins (Sonenberg and Hinnebusch, 2009). In particular, the recognition of the initiation codon of the canonical open reading frame (ORF) by ribosomes is crucial to obtaining functional proteins. While it is well-established that most translation starts at the AUG codon (Alberts et al., 2014; Krebs et al., 2018), AUG triplets can occur with an approximate frequency of every 4^3^ nucleotides (*i.e.*, 64 nt), presenting a serious challenge for efficient ribosomal recognition of the AUG codon corresponding to the canonical ORF in a given mRNA.

Eukaryotic cells are known to tackle the challenge by using a “scanning” mechanism (Kozak, 1978, 1989) based on the 43S preinitiation complex (PIC), comprised of a 40S ribosomal subunit, several eukaryotic initiation factors (eIFs), methionyl initiator transfer RNA (Met-tRNAi), and guanosine triphosphate (Hinnebusch, 2014; Merrick, 1992; Merrick and Pavitt, 2018; Pelletier and Sonenberg, 2019; Pestova and Kolupaeva, 2002). PIC scanning starts with attachment to the 5′-cap of an mRNA, after which the PIC migrates along the mRNA one nt at a time searching for the AUG codon: successive triplets enter the P-site of the 40S ribosomal subunit, where they are inspected for complementarity to the Met-tRNAi anticodon (Cigan et al., 1988; Jackson et al., 2010; Kozak, 1978, 1989; Saini et al., 2010).

According to current understanding, the PIC remains tethered to the eukaryotic mRNA (without jumping) during scanning (Hinnebusch, 2017; Kozak, 1978). This working model is supported by evidence showing that the most upstream (*i.e.*, 5′) AUG triplet is preferentially used as the primary initiation codon (Cigan et al., 1988; Kozak, 1984, 2002); insertion of an additional AUG triplet upstream (uAUG) of the annotated AUG triplet (aAUG, the initiation codon of the canonical ORF) can prevent translation initiation at the aAUG (Cuperus et al., 2017; Dvir et al., 2013; Hinnebusch, 2011; Johnstone et al., 2016; Kozak, 2002; Noderer et al., 2014; Zhang et al., 2019). Further, the insertion of a strong mRNA secondary structure between the 5′-cap and the aAUG can also prevent translation initiation (Berthelot et al., 2004).

Since the PIC starts scanning at the 5′-cap of an mRNA and the aAUG codon is somewhere downstream (*i.e.*, 3′), PIC scanning results in a net 5′–3′ ribosomal movement (Hinnebusch, 2014; Spirin, 2009). Currently, there are two competing models to explain the directionality of individual scanning steps (*i.e.*, one nt each step). The more established model that is commonly described in textbooks is the strictly unidirectional scanning model (Alberts et al., 2014; Kozak, 1984, 2002; Krebs et al., 2018), which posits that the PIC scans exclusively in the 5′–3′ direction (**Fig. 1A**). Sometimes the PIC misses an AUG triplet, an event termed “leaky scanning”, and will continue to scan the mRNA further downstream, which can enable access to downstream AUGs (dAUG) by the PIC. The scanning process proceeds until an AUG triplet is recognized (Hinnebusch, 2017; Jackson et al., 2010; Kozak, 2002).

**Fig. 1.**
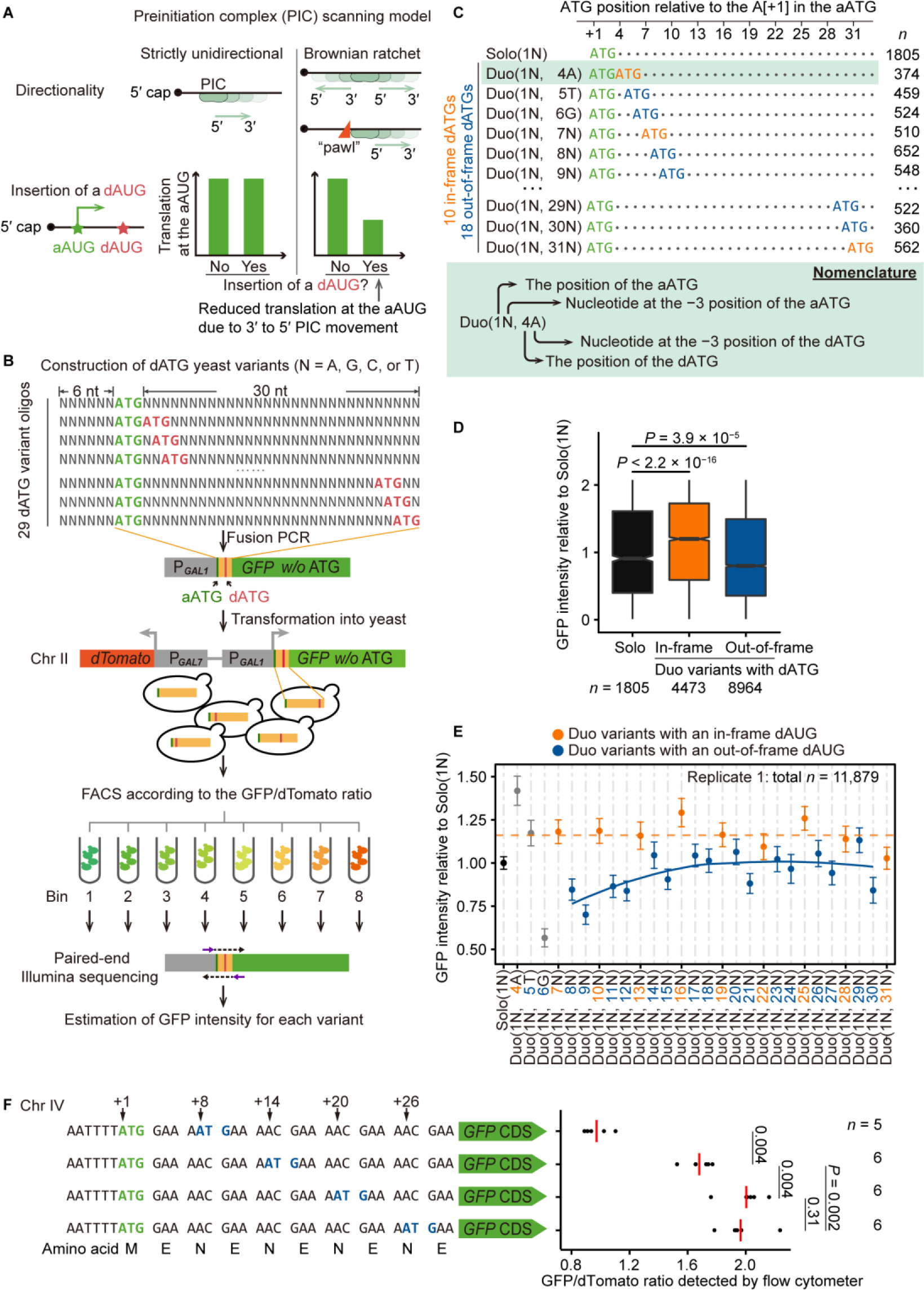
Testing PIC scanning models using thousands of dATG variants. (A) Predictions of the strictly unidirectional model and the Brownian ratchet scanning model on translational efficiency when an AUG is inserted downstream of the aAUG. (B) High-throughput construction of dATG variants with doped nucleotides and detection of GFP intensity *en masse* via FACS-seq. The aATGs of GFP are shown in green, and the dATGs are shown in red. (C) Nomenclature of Solo and Duo variants (grouped by the nucleotides at the −3 position of ATGs). The aATGs, in-frame dATGs, and out-of-frame dATGs are shown in green, orange, and blue, respectively. Each dot in the sequences represents a nucleotide that cannot form an ATG triplet or an in-frame stop codon. The number of variants (*n*, combined from two replicates) is shown on the right for each group of variants. (D) Boxplot shows GFP intensities (normalized by dTomato intensity, here and elsewhere in this study when presenting FACS-seq data) for Solo and Duo variants. *P* values were given by the Mann-Whitney *U* tests. (E) The average GFP intensities (dots) and the 95% confidence intervals (error bars) of Duo variants with the dATG at locations ranging from +4 to +31. The dashed line in orange represents the average GFP intensity of all Duo variants with an in-frame dATG, and the blue curve represents the local regression line generated by *R* function “geom_smooth” (span = 1) for the Duo variants with an out-of-frame dATG. Duo variants with the dATG at positions from +4 to +6 (shown in grey) had fixed nucleotides at the −3 position of the dATG (due to the aATG), and therefore, were not used to fit the local regression line. The plot shows the results for variants identified in biological replicate 1, and the results of biological replicate 2 are shown in Fig. S2. (F) A small-scale experiment that strictly controlled the flanking sequence of the dATG. The GFP/dTomato fluorescence ratio estimated for each replicate is shown in a dot and the average value is shown in the red line. *P* values were given by the Mann-Whitney *U* tests. See also Fig. S1, Fig. S2, and Table S5.

By contrast, the Brownian ratchet scanning model (Berthelot et al., 2004; Spirin, 2009) proposes that the PIC can migrate along an mRNA in both 5′–3′ and 3′–5′ directions governed by Brownian motion, as all movements of particles with similar size of ribosomes involve Brownian motion (Alekhina and Vassilenko, 2012; Spirin, 2009). The oscillation is directionally rectified through occasional placement of a “pawl” (*i.e.*, possibly RNA-binding proteins) on the mRNA at the trailing side of the PIC, restricting the 3′–5′ movement of the PIC beyond the pawl (**Fig. 1A**). In this model, even if an aAUG is missed by the PIC, it may be recognized in a second or subsequent scan as the PIC oscillates back and forth. A proximal dAUG, if present, can retain PICs that miss the aAUG, reducing the chance for a second (or more) inspection of the aAUG. In other words, the unidirectional scanning model relies on strictly sequential (5′–3′) decision-making involving more than one AUG in translation initiation, while in the Brownian ratchet model initiation decisions are competitive rather than strictly sequential for closely spaced AUGs.

The fundamental difference between the two models is whether the PIC can move in the 3′–5′ direction, which can be experimentally determined by insertion of out-of-frame dATGs. The strictly unidirectional scanning model predicts that dATGs will have no effect on translation initiation of the canonical ORF, whereas the Brownian ratchet scanning model predicts that a dAUG can exhibit an inhibitory effect on initiation at the aAUG due to competition as the translation initiation site (**Fig. 1A**). Consistent with the strictly unidirectional model, previous work has shown that the first AUG codon can exclusively serve as the site of translation initiation even when a second AUG is located within just a few nt downstream (Kozak, 1995). However, other studies with genetically modified overlapping bicistronic mRNAs from *Turnip yellow mosaic virus* and *Influenza virus B* (Matsuda and Dreher, 2006; Williams and Lamb, 1989) have revealed that the initiation frequency of an upstream ORF can be reduced by the presence of a proximal, downstream, overlapping ORF, thus implying the presence of 3′–5′ PIC scanning (Hinnebusch, 2014; Jackson et al., 2010; Matsuda and Dreher, 2006). These findings indeed raised many questions, and led us to investigate whether such 3′–5′ scanning observed in bicistronic viral mRNAs could also occur endogenously in monocistronic eukaryotic mRNAs.

Here, to compare the strictly unidirectional vs. Brownian ratchet scanning models, we generated thousands of green fluorescent protein (*GFP*) reporter gene sequence variants, each containing an out-of-frame ATG downstream of the ATG corresponding to the canonical ORF. We measured the fluorescence intensity of each variant, then performed computational simulations and estimated the leakage rate of each scan, as well as the number of scans for each triplet, before the ribosome eventually migrated to downstream regions of the mRNA. Our results reveal several general rules governing ribosomal scanning, ultimately supporting a Brownian ratchet model. These findings also enhance our understanding of how point mutations that introduce dATGs can lead to dysregulation of gene expression that is associated with human diseases.

## Results

### Generation of thousands of dATG yeast variants

Previous studies have widely observed reduced protein abundance upon the addition of an out-of-frame uATG (Cuperus et al., 2017; Dvir et al., 2013; Noderer et al., 2014), indicating that PICs scan continuously along the mRNA in the 5′–3′ direction (Kozak, 1978, 1989, 2002). By the same logic, a reduction in GFP intensity following the addition of an out-of-frame dATG will indicate that ribosomes can also scan in the 3′–5′ direction (**Fig. 1A**). In this study, we investigated the occurrence and prevalence of 3′–5′ PIC movement by inserting ATGs downstream of the aATG of a GFP reporter and then detecting the impacts on GFP synthesis (*i.e.*, through differences in GFP intensity). To avoid reaching conclusions that are caused by some specific flanking sequences (*i.e.*, confounding factors), as in viruses that may use specific sequences to regulate PIC scanning for overlapping ORFs in their bicistronic mRNAs (Matsuda and Dreher, 2006; Williams and Lamb, 1989), we generated a large number of sequence variants, each with a dATG inserted in various sequence contexts (**Fig. 1B**).

Specifically, we introduced dATGs by chemically synthesizing a 39-nt DNA oligo, with six upstream and thirty downstream doped (*i.e.*, random) nucleotides (N = 25% A + 25% T + 25% G + 25% C) around a fixed ATG triplet (designated as the aATG, **Fig. 1B** **and Fig. S1A**). ATG triplets (either in-frame or out-of-frame) could then form randomly within the 30-nt downstream region of each individual construct, ultimately resulting in a representative library containing dATGs at each successive position downstream of the aATG in various sequence contexts. To increase the fraction of dATG-containing variants, we further synthesized 28 additional DNA oligos, each with a dATG fixed at one of the 28 possible downstream triplet positions (**Fig. 1B** **and Fig. S1A**). We fused these DNA oligos with the full-length *GFP* sequence (with its initiation codon omitted, **Fig. S1A**), and integrated the fusion constructs individually into the same locus in Chromosome II of the yeast genome. We also inserted *dTomato*, encoding a red fluorescent protein, into a nearby genomic region to normalize GFP intensity (**Fig. 1B** **and Fig. S1A**).

We measured the GFP intensities *en masse* through fluorescence-activated cell sorting (FACS)-seq for individual variants, as described in a previous study (Chen et al., 2017). Briefly, we sorted yeast cells into eight bins according to GFP intensity (here and elsewhere in this study, normalized by dTomato intensity). Based on the variant frequencies in high-throughput sequencing reads of the eight bins, and the median GFP intensity and the number of cells belonging to each bin, we calculated the GFP intensity for each variant as the weighted average GFP intensity across the eight bins (**Fig. S1A**).

To verify the accuracy of GFP intensities measured *en masse*, we randomly isolated 20 clones from the yeast library, and individually measured their GFP intensities by flow cytometer. There was good consistency between the GFP intensities measured *en masse* and individually (Pearson’s correlation coefficient *r* = 0.99, *P* = 1 × 10^−19^, **Fig. S1B**). We measured GFP intensity of the yeast variants in two biological replicates, and the values were highly correlated for 18,950 variants shared between both experiments (Pearson’s correlation coefficient *r* = 0.99, *P* < 2.2 × 10^−16^, **Fig. S1C**). Consequently, we pooled dATG variants from both replicates in subsequent data analyses (**Fig. S1D,** the average GFP intensity was used for variants shared by the two replicates), if not otherwise specified.

We performed two positive control analyses to examine the data quality. First, the GFP intensity of variants with in-frame stop codons formed in the 30-nt region downstream of aATG was lower than that of variants without in-frame stop codons (*P* < 2.2 × 10^−16^, Mann-Whitney *U* test; **Fig. S1E**). Second, the variants containing in-frame uATGs showed elevated GFP intensity compared to variants without uATGs (*P* < 2.2 × 10^−16^, Mann-Whitney *U* test; **Fig. S1F**), most likely because the second in-frame AUGs could function as an auxiliary initiation site for GFP translation. In contrast, the variants containing out-of-frame uATGs showed reduced GFP intensity (*P* < 2.2 × 10^−16^, Mann-Whitney *U* test; **Fig. S1F**), likely because they can prevent translation in the reading frame of *GFP*. These observations bolstered our confidence to compare GFP intensities among the dATG variants in our study. Note that we excluded the variants containing in-frame stop codons or uATGs from the subsequent analyses, to avoid their potential impacts on GFP intensity.

### The nucleotide at the −3 position affects the leakage rate at the AUG

Seminal studies analyzing the consensus sequence across genes (Hamilton et al., 1987; Kozak, 1986, 1987, 2002) led to the hypothesis that some flanking sequences could facilitate translation initiation (known as the Kozak sequence). To determine if the sequences flanking the aAUGs exerted any detectable influence on the GFP intensities measured in our yeast library, we grouped the 21,598 variants according to the nucleotide type at each position, and estimated the average GFP intensity for each of the four variant groups at each position (**Fig. S2A**). Briefly, placing different nucleotides at the −3 position (relative to the A[+1] in the aATG codon) led to the highest variation in GFP intensity compared to variation related to different nucleotides at other positions (from −6 to +15, **Fig. S2A**). At the −3 position, “A” conferred the highest GFP intensity, followed by G, C, and finally T. This observation is qualitatively consistent with the prevalence of A at the −3 position among the 500 most highly expressed genes in the yeast genome (**Fig. S2B**). For simplicity, we hereafter refer to the ATG context using the nucleotide at the −3 position; in the order from “strong” to “weak” are the A-, G-, C-, and T-contexts.

We hypothesized that the observed differences in the strength of the sequence context could be directly related to leaky scanning. To gauge the leakage rate in endogenous yeast genes, we retrieved the mRNA sequences protected by small ribosomal subunits (SSU footprints) during RNase I digestion obtained in a previous study (Archer et al., 2016). We counted the sequencing reads 40-nt upstream and 40-nt downstream of the aAUG for each gene, and defined the leakage rate of a gene as the fraction of downstream footprints among all footprints within the 80-nt range (**Fig. S2C**). We found that the −3 position had the largest variation in the leakage rate among the four nucleotide types across the flanking sequences (from −6 to +6, **Fig. S2C**), and that the results remained qualitatively unchanged with a larger window size (80-nt upstream and downstream, **Fig. S2C**). The GFP intensity (**Fig. S2A**) was in the reverse order as the leakage rate across the four nucleotides at the −3 position, indicating a tight association between leaky scanning and the strength of sequence context.

### Frame- and distance-dependent translational inhibition by dAUGs

Prior to measuring the effects of dAUGs on GFP intensity, we considered the variation in the number, position, and context of ATGs among the variants in the yeast library, to establish a standardized and clear nomenclature for these variants. Some variants had only one ATG in the 39-nt region (*i.e.*, the designed aATG), and were therefore denoted as “Solo” variants. Some variants had one additional ATG in the 30-nt downstream region (*i.e.*, the dATG) and were thus designated as “Duo” variants. In addition, the names of variants include the position and context of the aATG and dATG (if present). For example, Duo(1N, 4A) represents variants with two ATGs: the aATG having any one of the four nucleotides (N) at the −3 position and a dATG at the +4 position with an A in its −3 position (**Fig. 1C**). We subsequently focused on analysis of 1805 Solo and 13,437 Duo variants.

In our design, dATGs were introduced at a total of 28 positions, among which 10 were in-frame and 18 out-of-frame, relative to the *GFP* reading frame (**Fig. 1C** **and Fig. S1A**). To investigate whether out-of-frame dAUGs can inhibit translation initiation from the aAUG, we grouped the Duo variants according to the reading frames of their dATGs. The results showed that the Duo variants containing in-frame dATGs showed elevated GFP intensity compared to Solo variants (*P* < 2.2 × 10^−16^, Mann-Whitney *U* test; **Fig. 1D**), as variants containing in-frame uATGs (**Fig. S1F**). In sharp contrast, Duo variants harboring an out-of-frame dATG showed reduced GFP intensity compared to Solo variants (*P* = 3.9 × 10^−5^, Mann-Whitney *U* test, **Fig. 1D**), strongly suggesting that out-of-frame dAUGs can inhibit translation initiation at the aAUG in a translation-dependent manner. The fraction of reduction in GFP intensity for Duo variants relative to Solo variants is termed as the “inhibitory effect” subsequently.

To test if these inhibitory effects of out-of-frame dAUGs were dependent on the distance between aATG and dATG, we grouped the Duo variants according to the position of their dATG and then estimated the average GFP intensity for each group. The inhibitory effect gradually declined with increasing aATG-dATG distance (**Fig. 1E** and **Fig. S2D**), and no inhibitory effects were evident at aATG-dATG distances of ∼17 nt or greater (**Fig. 1E** and **Fig. S2D**). These observations indicated that translation initiation decisions involving two proximal, potential AUGs were not strictly sequential, but competitive. Note that the placement of dATGs at various positions did not significantly alter the synonymous codon usage or the formation of mRNA secondary structure in the 30-nt variable sequence downstream of the aAUG (**Fig. S3**), two factors known to affect translation elongation, and therefore, protein synthesis (Chu et al., 2014; Kudla et al., 2009; Yang et al., 2014).

We then performed an additional, small-scale experiment that strictly controlled the flanking sequence to further characterize the distance-dependent inhibitory effect of dAUGs. Specifically, we introduced an out-of-frame dATG at +8, +14, +20, or +26 positions downstream of the aATG (**Fig. 1F**). To exclude any potential impacts of the peptide sequence on GFP intensity, we used only synonymous mutations to introduce these out-of-frame dATGs. The results showed that proximal out-of-frame dATGs indeed reduced GFP intensity, while increases in distance between the two ATGs resulted in a gradual increase in GFP intensity. Beyond 20 nt, the negative impacts on translation initiation were no longer detectable (**Fig. 1F**). Collectively, these results established that out-of-frame dATGs could inhibit GFP synthesis, and that these inhibitory effects decreased with increasing distance from the aATG.

### Context-dependent translational inhibition by dAUGs

The frame- and distance-dependent inhibitory effects of dAUG suggested that ribosomes could sometimes scan in the 3′–5′ direction, which was compatible with the Brownian ratchet scanning process wherein PICs oscillate in both 5′–3′ and 3′–5′ directions, scanning each successive triplet multiple times. An aAUG that is not recognized by the PIC in the first scan may be recognized in a subsequent scan. When a dAUG is inserted near the aAUG, a PIC that misses the aAUG may be instead retained by that nearby dAUG if it is recognized, thereby reducing the likelihood that a PIC will oscillate 3′–5′ and recognize the aAUG. As the aAUG–dAUG distance increases, there is an increased probability that a given PIC will turn to the 3′–5′ direction before it encounters a dAUG (**Fig. S4A**).

The Brownian ratchet scanning model further predicted that the aAUG–dAUG competition was dependent on the leaky scanning at the aAUG. To test if the observed inhibitory effect of proximal out-of-frame dAUGs is indeed related to the leaky scanning at the aAUG, we divided the Duo variants into four groups based on their aATG −3 context. We found that the inhibitory effect of dATGs was greater when the aATG was in a weaker context (*i.e.*, higher leakage rate, **Fig. 2A** and **Fig. S2E**), which indicated that leaky scanning at aAUGs contributed to dAUG inhibition of translation initiation. To then determine whether these inhibitory effects were due to translation initiation at the dAUG, we also divided the Duo variants into four groups according to their dATG −3 context. We found that the inhibitory effect was greater when the dATG was in a stronger context, indicating the competition of translation initiation between the two AUGs (**Fig. 2B** and **Fig. S2E**).

**Fig. 2.**
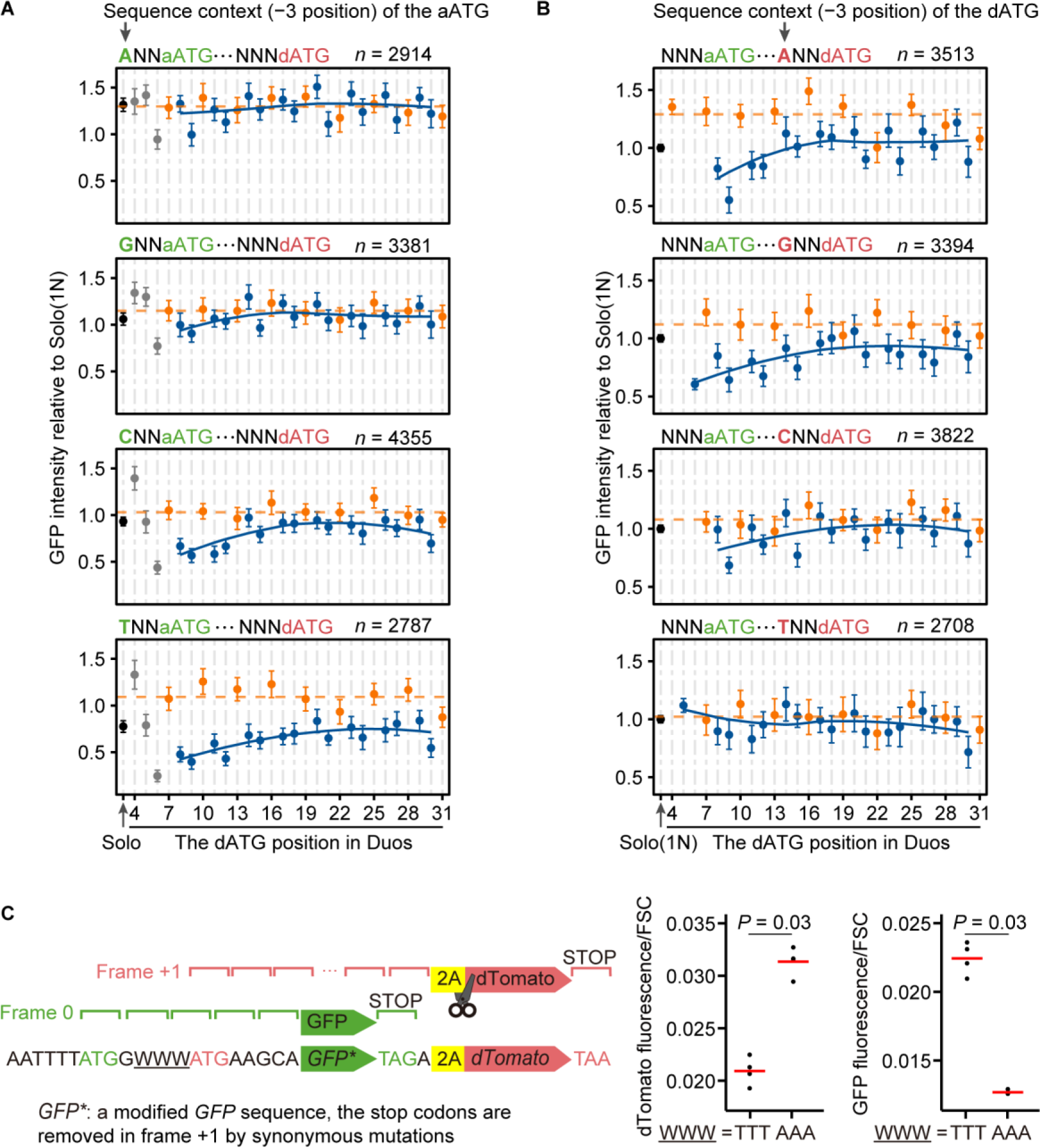
Context-dependent inhibitory effects on protein synthesis by proximal out-of-frame dATGs. (A-B) The average GFP intensities (dots) and the 95% confidence intervals (error bars) of Duo variants, grouped according to sequence contexts of the aATG or dATG (letters in green or red, respectively). The numbers of A-, G-, C-, and T-context Solo variants are 341, 383, 719, and 362, respectively. (C) The overlapping dual-fluorescence reporter experiment to confirm the competition between aAUG and dAUG as the translation initiation site. The reporter is composed of a modified *GFP* gene in frame 0, sequences encoding a 2A self-cleaving peptide in frame +1, and a *dTomato* gene in frame +1. The GFP fluorescence was normalized by the measurement of forward scatter (FSC), which measures the cell size. The values of individual replicates are shown by black dots (*n* = 4) and the mean is shown by the red line. *P* values were given by the Mann-Whitney *U* tests. See also Figs. S2 and S3.

To confirm this apparent competition between aAUGs and dAUGs for translation initiation, we performed an experiment using an overlapping dual-fluorescence reporter in which GFP was translated from an aAUG in a weak context, and dTomato was translated from a proximal out-of-frame dAUG (+8 position, **Fig. 2C**). Placing the dATG in two different contexts, we measured both green and red fluorescence intensities with flow cytometry. We observed that dTomato intensity increased with increasing strength of dATG context (*i.e.*, lower leakage rate) while GFP intensity was substantially reduced (**Fig. 2C**). These results confirmed that translation initiation decisions between two closely spaced AUGs were determined in a competitive manner.

### Proximal out-of-frame dAUGs lead to reduced mRNA levels via nonsense-mediated mRNA decay (NMD)

Our findings above thus suggested that proximal out-of-frame dAUGs could compete with aAUG for translation initiation. Since out-of-frame termination codons are abundant in the *GFP* coding sequence (see Methods), we predicted that if translation indeed initiated at a proximal out-of-frame dAUG, a long distance should remain between its (also out-of-frame) termination codon and the poly(A) tail, a signal for mRNA degradation by the NMD pathway (Kurosaki et al., 2019; Losson and Lacroute, 1979; Muhlrad and Parker, 1999). To test if the insertion of proximal out-of-frame dAUGs can result in lower *GFP* mRNA stability, we measured the mRNA levels *en masse* for each variant in the library, as described in previous work (Chen et al., 2017). Briefly, we used Illumina sequencing to determine the mRNA levels of each variant, which was normalized by the number of cells for each variant (**Fig. 3A**). Since the mRNA levels of dATG variants were highly correlated between two biological replicates (Pearson’s correlation coefficient *r* = 0.86, *P* < 2.2 × 10^−16^, **Fig. S5A**), we pooled dATG variants from both replicates in subsequent data analyses. We grouped the Duo variants according to the position of their dATGs, as well as by the aATG and dATG contexts. The results showed that variants with an out-of-frame dATG reduced mRNA levels in a distance-dependent manner, particularly when the aATG resided in a weaker context (**Fig. S5B**) and dATG resided in a stronger context (**Fig. 3B**), indicating that the initiation competition from proximal out-of-frame dAUGs could reduce the mRNA level.

**Fig. 3.**
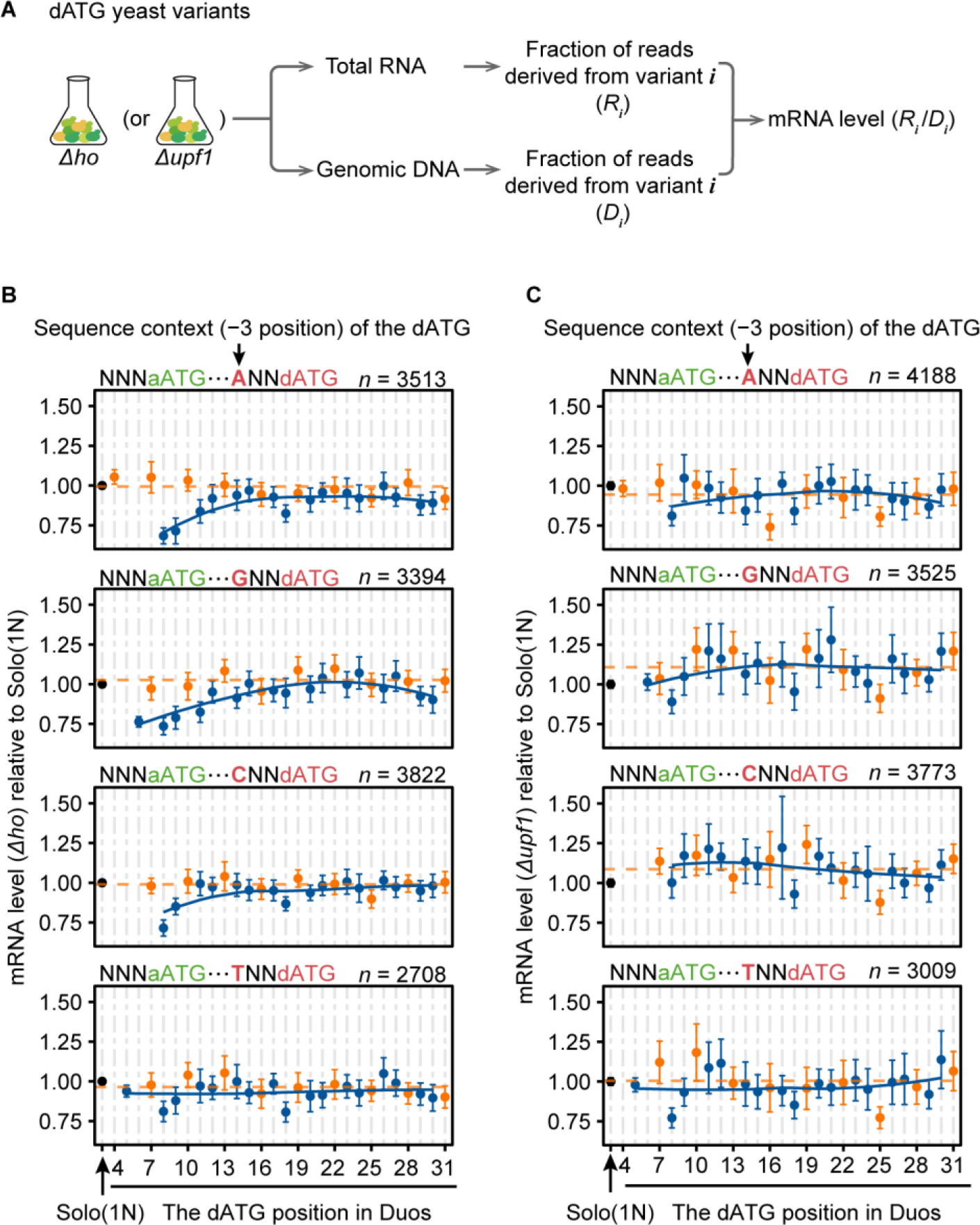
The reduction in the mRNA level caused by proximal out-of-frame dATGs, via the NMD pathway. (A) The experimental procedure for high-throughput determination of the mRNA levels for individual dATG variants. (B–C) The average mRNA levels (dots) and the 95% confidence intervals (error bars) of Duo variants, in the background of *Δho* (B) and *Δupf1* (C), grouped according to sequence contexts of the dATG (letters in red). The numbers of N-context Solo variants are 1805 (*Δho*) and 2194 (*Δupf1*).

To then determine whether the reduction in the mRNA level was caused by the NMD pathway, we knocked out *UPF1*, the gene encoding an RNA helicase required for initiating NMD in eukaryotes (He et al., 2003; Leeds et al., 1991), and created a new yeast library containing a total of 41,456 variants in the background of *Δupf1* (**Fig. 3A** and **Fig. S5A**). We measured the mRNA levels of these variants and found that the reduction in mRNA levels we previously observed in Duo variants with proximal out-of-frame dAUGs was abolished in the absence of *UPF1* (**Fig. 3C** and **Fig. S5C**). These observations indicated that the NMD pathway activated by translation initiation at out-of-frame dAUGs could exacerbate the inhibitory effect of proximal dAUGs at the translational level.

To exclude the possibility that the distance-dependent inhibitory effect of out-of-frame dATGs is associated with variation in the activation efficiency for NMD, which was reported depending on the position of their corresponding stop codons (Muhlrad and Parker, 1999), we further excluded 1678 variants that contained out-of-frame stop codons in the variable region in the same reading frame of the dATG. Therefore, all 3613 (or 3673) Duo variants containing frame +1 (or +2) dAUG would terminate translation at a fixed location further downstream (+56 or +60, see Methods). The inhibitory effect of proximal out-of-frame dATGs remained observed (**Fig. S5D**), excluding the variation in NMD efficiency among dATG variants as a confounding factor.

### Distance-dependent translational inhibition by upstream AUGs (uAUGs)

In general, proximity to the 5′-cap grants an AUG triplet some advantage for initiating translation since they are scanned first (Hinnebusch, 2014; Kozak, 2002). Consistent with this hypothesis, it was widely reported that out-of-frame uAUGs could inhibit translation at the aAUG because the uAUG can retain a proportion of PICs that would otherwise initiate translation at the aAUG (Dvir et al., 2013; Johnstone et al., 2016; Kozak, 2002). Given our results showing competition for initiation between a closely spaced aAUG-dAUG pair, we further predicted that a closely spaced uAUG-aAUG pair would also compete for translation initiation. That is, when a uAUG is near the aAUG, a PIC that misses the uAUG (due to leaky scanning) may be retained by the nearby aAUG, thereby reducing the likelihood that the PIC will oscillate 3′–5′ and recognize the uAUG. Therefore, the Brownian ratchet scanning model further predicted that the inhibitory effect by an out-of-frame uAUG should diminish with decreasing uAUG-aAUG distance (**Fig. 4A**).

**Fig. 4.**
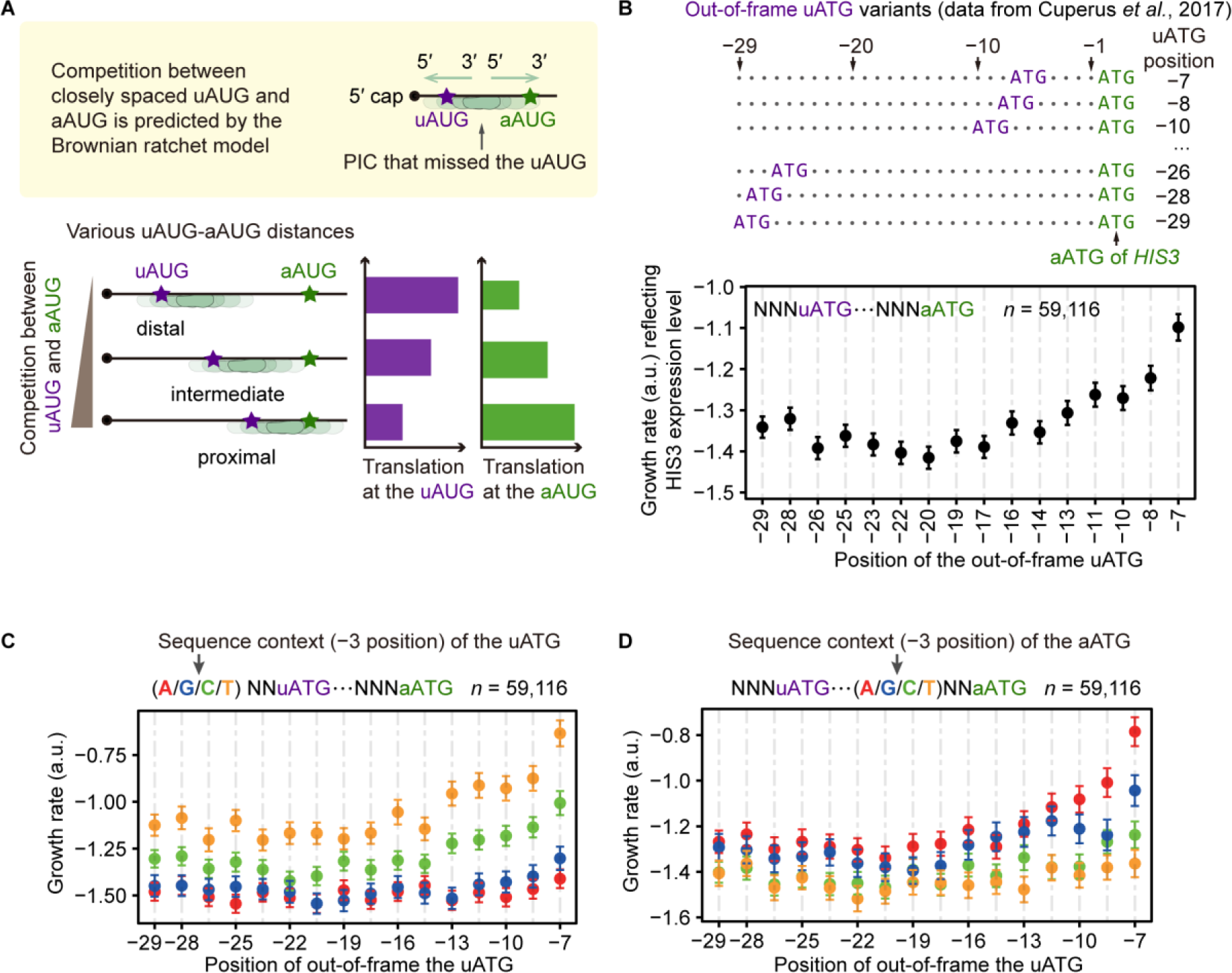
The distance-dependent inhibitory effect of out-of-frame uAUGs on translation initiation at the aAUG. (A) Predictions of the Brownian ratchet scanning model on translational efficiency at the aAUG, when an out-of-frame AUG is introduced upstream at various positions. (B) The average growth rates (dots) and the 95% confidence intervals (error bars) of uATG variants, grouped by the positions of their uATGs relative to the aATG. In Cuperus *et al*. (2017), ATGs were inserted upstream of the aATG of *HIS3*, a gene that encodes an enzyme in the histidine biosynthesis pathway; consequently, the growth rate of a variant in the medium without histidine reflected the protein abundance of HIS3. (C–D) The growth rates of uATG variants, grouped by the nucleotide at the −3 position of the uATG (C) or aATG (D).

To test if the inhibitory effect of an out-of-frame uAUG indeed depends on its distance to the aAUG, we retrieved previously reported data from a study that generated a large number of variants, each with an ATG randomly inserted at various positions upstream of the *HIS3* aATG (Cuperus et al., 2017). In the experiment, yeast cells were cultured in the absence of histidine, and therefore, the growth rate measured by the authors qualitatively reflected the protein abundance of HIS3. We grouped these variants according to the positions of the inserted out-of-frame uATGs, similar to our grouping of dATGs in our library. The results showed that growth rates increased with decreasing uATG-aATG distance, a trend which was only detectable for uATGs within the ∼17 nt upstream of the aATG (**Fig. 4B**). We also observed that the inhibitory effect of a proximal, out-of-frame uATG was weaker in variants with the uATG in a weaker context (**Fig. 4C**) or with the aATG in a stronger context (**Fig. 4D**). Taken together, these results showing distance- and context-dependent inhibitory effects by out-of-frame uATGs suggested translational competition between aAUG and a proximal uAUG.

### Each successive triplet is scanned by the PIC approximately 10 times

The translational competition we detected between closely spaced AUGs was qualitatively consistent with the Brownian ratchet scanning model in which the PIC is tethered to mRNA and progresses toward 3′-end under a Brownian ratchet mechanism. Notably, the quantification of GFP intensity we conducted for thousands of variants in this study provided us with an opportunity to estimate the parameters of PIC scanning, such as the number of scans for each triplet, the frequency that a pawl (*i.e.*, the 5′-block) is placed along the mRNA, and the efficiency of AUG recognition by the PIC. To quantitatively investigate the PIC scanning process, we simulated the Brownian ratchet scanning process using a modified random walk model, as the PIC movement consists of a succession of random steps on the discrete positions along the “one-dimensional space” of a linear mRNA (**Fig 5A**).

**Fig. 5.**
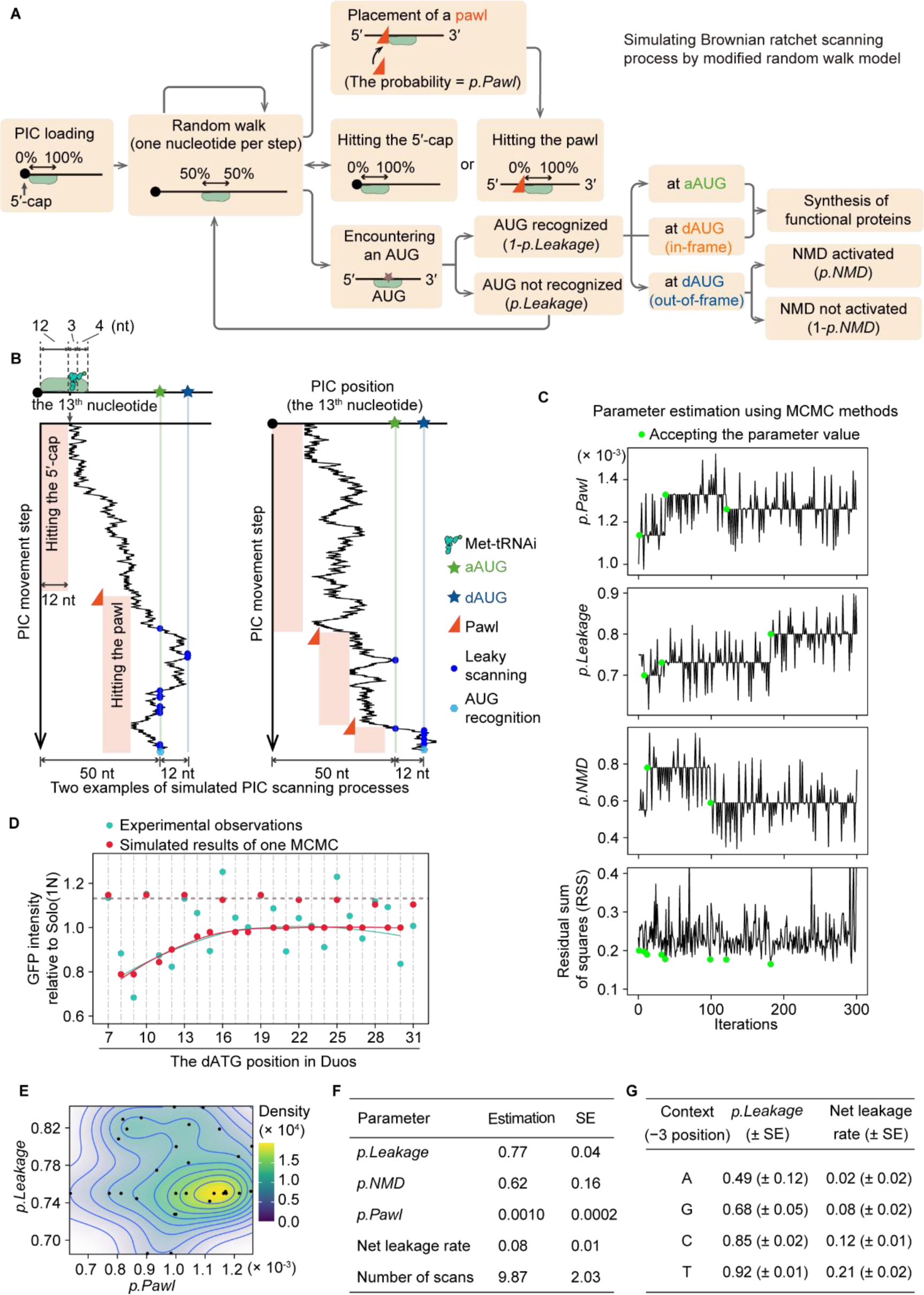
Estimation of parameters in the Brownian ratchet scanning model by MCMC algorithms. (A) A flowchart illustrating decisions involving PIC movement, translation initiation (or leakage), placement of the “pawl”, and the activating of the NMD pathway. (B) Two trajectories of the PIC exemplify the simulated PIC scanning process along the mRNA. According to Archer *et al*. (2016), the PIC covers 19 nucleotides of an mRNA, where the 13^th^–15^th^ nucleotides are inspected for complementarity to the Met-tRNAi anticodon. The position of the 13^th^ nucleotide is used to denote the PIC position. (C) Trace plots show changes in values of *p.Pawl*, *p.Leakage*, and *p.NMD* in an MCMC chain. Green dots mark the MCMC iterations that accepted a new parameter value due to a reduction in the RSS. (D) The comparison of the observed GFP intensities of Duo variants in the yeast experiments (two replicates combined) and the simulated GFP intensities using one of the 30 sets of optimized parameters by the MCMC algorithms. The results of all 30 sets of optimized parameters are shown in Fig. S7. (E) Two-dimensional density plot shows the distribution of the outcome values of *p.Pawl* and *p.Leakage* among the 30 MCMC chains (shown by dots). Two additional density plots involving *p.NMD* are shown in Fig. S6C. (F–G) Estimated parameters and standard error (SE) in the Brownian ratchet scanning model. See also Figs. S6 and S7.

We specified the following parameters in our random walk model. The PIC occupies 19 nt of the mRNA during scanning (Archer et al., 2016), and the triplet at the 13^th^–15^th^ position is the P-site of the PIC, where the inspection for complementarity to the Met-tRNAi anticodon occurs. The PIC started out at the 5′-cap and took one nt per step in either the 5′–3′ or 3′–5′ direction, with equal probability (*i.e.*, 50% each). However, the PIC could not move further upstream if its 5′-trailing side hits the 5′-cap or a pawl. The pawl was stochastically placed along the mRNA at the 5′-trailing side of a PIC (depending on the PIC location at the time) with the probability *p.Pawl* (**Fig 5A**). When an AUG enters the P-site of the PIC, in a probability of *p.Leakage* the AUG might not be recognized by the PIC, and in a probability of (1 − *p.Leakage*) the AUG was recognized by the PIC and started translation. Note that in our model AUG triplets could be recognized in either 5′–3′ or 3′–5′ PIC movement. Considering that the NMD pathway can reduce mRNA levels, and consequently protein abundance, we used the parameter *p.NMD* to determine the probability of activating NMD when an out-of-frame dAUG is recognized by the PIC (**Fig 5A**).

We employed a Markov Chain Monte Carlo (MCMC) algorithm (Hastings, 1970; Metropolis et al., 1953) to calculate numerical approximations for the probability parameters in the Brownian ratchet scanning model. To compare with experimental measurements of GFP intensity (**Fig. 1E**), we simulated the Brownian ratchet scanning process for 25 Duo variants with N-context dAUGs (representing an “average” dAUG) positioned between +7 and +31. To explore the relevant parameter space by the MCMC sampler, we first tested 1000 parameter sets for *p.Pawl*, *p.Leakage*, and *p.NMD* (each with 10 values ranging from 0.001 to 0.8, **Fig. S6A**). For each parameter set, we generated 100 simulations of the PIC scanning process for each Duo variant (two examples are shown in **Fig. 5B**) and estimated GFP intensity based on the number of simulations in which translation was initiated at the aAUG. We identified the 10 parameter sets that showed the smallest residual sum of squares (RSS) for the 25 Duo variants, and initiated the MCMC simulation using the median value for each parameter among the 10 parameter sets: *p.Pawl* = 0.001, *p.Leakage* = 0.75, and *p.NMD* = 0.55 (**Fig. S6B**).

We then ran the MCMC algorithm for 300 iterations, in which we sequentially replaced each of the three probability parameters with a random number generated from a uniform distribution (see Methods). For each iteration, we calculated the RSS from the simulated and observed GFP intensities in our yeast library, and used the RSS as a proxy to optimize the parameters. If the RSS decreased, the previous set of parameters was replaced by the new parameters, whereas if the RSS increased, the previous set of parameters remained unchanged (**Fig. 5C**).

The parameters reached the stationary distribution at the end of 300 iterations (**Fig. 5C– D**), and the GFP levels observed in the experiments were largely recapitulated by our simulated ratchet and pawl model of PIC scanning (**Fig. 5D**). Specifically, the inhibitory effects of dAUG were detected in the simulation, and became undetectable beyond ∼17 nt. To obtain a reliable estimation of the parameters, we repeated 30 times of MCMC, and found that the parameters were stable after 300 iterations in all MCMCs (**Fig. 5E**, **Fig. S6C**, and **Fig. S7**). The average parameter values that resulted in the smallest RSS were as follows: the probability of adding a pawl to the mRNA was ∼ 1 out of 1000 PIC steps; the average leakage rate for each single scan was 77%; and the NMD rate was 62% (**Fig. 5F**). Based on these values, we estimated that, on average, each triplet was scanned ∼10 times by the PIC before a pawl was placed between the triplet and the PIC, resulting in a net leakage rate of 8% (*i.e.*, on average 8% of PICs eventually miss an AUG after multiple scans).

So far, we simulated using an N-context dAUG and neglected any difference in the leakage rate among sequence contexts of ATG triplets. To individually estimate *p.Leakage* for ATGs in the A-, G-, C-, or T-contexts, we fixed the values of *p.Pawl* (= 0.001) and *p.NMD* (= 0.62) and optimized context-specific *p.Leakage* by running the MCMC algorithm for another 100 iterations, based on the RSS estimated from the GFP intensities of variants Solo(1A), Solo(1G), Solo(1C), and Solo(1T) observed in our experiments (**Fig. 2A**). The average *p.Leakage* values among 30 times of MCMCs were 0.49, 0.68, 0.85, and 0.92 (**Fig. 5G** **and Fig. S6D**), corresponding to the net leakage rates of 0.02, 0.08, 0.12, and 0.21, for ATGs in the A-, G-, C-, or T-context, respectively (**Fig. 5G**). These numbers were qualitatively consistent with the leakage rates estimated from SSU footprints (**Fig. S2C**) and the nucleotide usage frequency observed in the yeast genome (**Fig. S2B**).

### Depletion of proximal out-of-frame dATGs in yeast and human genomes

Given the reduced efficiency for translation of canonical ORFs and possibility of enhanced synthesis of potentially cytotoxic peptides, we predicted that proximal out-of-frame dATGs would be generally deleterious. Therefore, we sought to test if proximal out-of-frame dATGs have been purged from the yeast genome by purifying selection. To this end, we counted the number of genes with ATGs at various positions downstream of the aATG across the yeast genome (**Fig. 6A**). The results showed that the number of out-of-frame dATGs increased gradually with distance from the aATG, and plateaued at around positions +20 and +21. The trend was statistically significant only in frame +1, probably because ∼80% of out-of-frame dATGs are located in frame +1 due to the preferred usage of some amino acids or codons. Moreover, the increase in the number of frame +1 dATGs was particularly apparent for dATGs in the A/G-context (**Fig. 6B**). Collectively, the scarcity of proximal, frame +1 dATGs, especially those in the stronger contexts, suggested the effects of purifying selection against these dATGs.

**Fig. 6.**
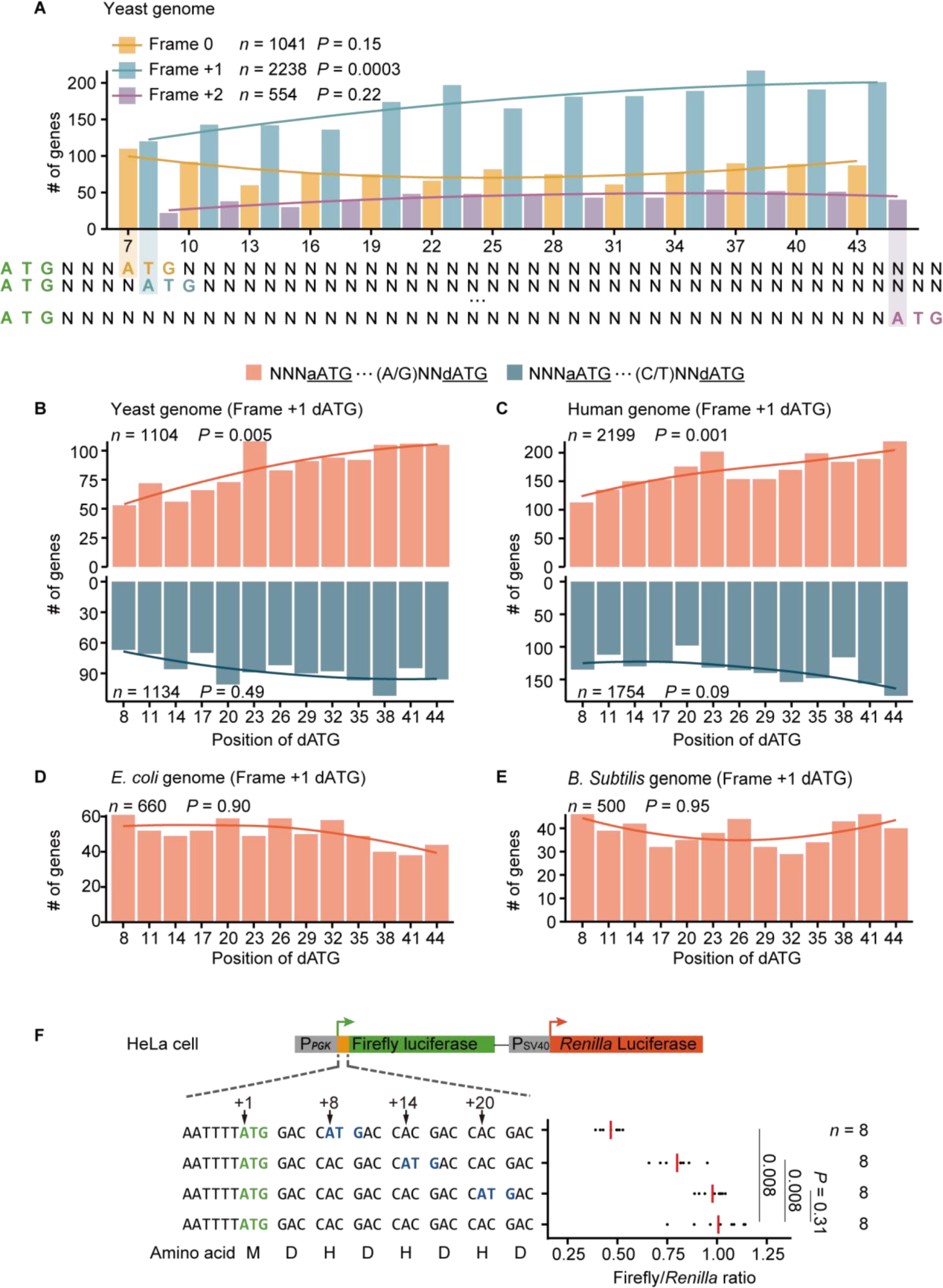
The depletion of proximal out-of-frame dATGs in eukaryotic genomes. (A) Meta-gene analysis shows the number of genes that harbor proximal dATGs at individual positions in the yeast genome. The curves represent the local regression line (span = 2). *P* values were given by the *G*-tests for uniform distribution. (B–E) Numbers of genes that harbor frame +1 dATGs at individual positions in the genomes of yeast (B), humans (C), *Escherichia coli* (D), and *Bacillus Subtilis* (E). The colors of the bars show the dATG context. *E. coli* and *B. Subtilis* are used as negative controls as no ribosomal scanning mechanism is required for translation initiation in prokaryotes. (F) The dual-luciferase reporter experiment to test the distance-dependent inhibitory effect of dAUGs in HeLa cells. The values of individual replicates (*n* = 8) are shown by black dots and the average values are shown by red lines. *P* values were given by the Mann-Whitney *U* tests.

To test if proximal out-of-frame dATGs were also purged from other eukaryotic genomes, we similarly counted the number of frame +1 dATGs at various positions in the human genomes. Similar to our analysis of the yeast genome, we found that the number of frame +1 dATGs increased with distance from the aATG, consistent with the observation in a previous study (Kochetov, 2005). And the trend was more obvious for dATGs in a stronger context (**Fig. 6C**). As a negative control, prokaryotes, which do not use the scanning mechanism to search the initiation codon (Kapp and Lorsch, 2004; Kozak, 1999), did not show the depletion of proximal out-of-frame dATGs (**Fig. 6D–E**). These observations implied that the Brownian ratchet scanning process likely generally affected the evolution of eukaryotic genomes by purging proximal out-of-frame dATGs.

Although the translation machinery of yeast and humans is largely identical, some differences have been reported in the components of eIFs (Hinnebusch, 2014). To test whether proximal out-of-frame dAUGs can indeed inhibit translation initiation at the aAUG in humans, we constructed a firefly and *Renilla* dual-luciferase reporter system and designed three additional variants, each with a frame +1 ATG introduced at a different location (+8, +14, and +20) downstream of the firefly luciferase aATG, using synonymous mutations (**Fig. 6F**). We transfected the reporters individually into HeLa cells and measured the *Renilla*-normalized firefly luciferase activity. In agreement with our findings in yeast, the results showed that proximal out-of-frame dATGs reduced firefly luciferase activity. Moreover, increases in the distance between the two ATGs resulted in a gradual increase in firefly luciferase activity, which was no longer distinguishable from the wild type in ATG variants located 20-nt downstream of the aATG (**Fig. 6F**). These results confirmed evolutionary conservation of the inhibitory effects on protein synthesis by proximal out-of-frame dATGs between humans and yeast.

### Cancer-related somatic mutations that introduce proximal out-of-frame dATGs are associated with reduced mRNA levels

Considering that cancer cells often accumulate numerous somatic mutations, it was reasonable to speculate that point mutations in these cells could introduce dATGs that reduced the translation initiation at the aAUG. Provided that aberrant translation initiation at out-of-frame dAUGs would result in reduced mRNA levels via NMD (**Fig. 3**), we used mRNA levels as a proxy to infer the inhibitory effects of dAUGs on aAUG translation. We retrieved transcriptome data collected from a total of 9418 cancer samples spanning 33 cancer types, as well as the somatic point mutations identified in these samples, from The Cancer Genome Atlas (TCGA) (Cancer Genome Atlas Research Network et al., 2013). To exclude potential interference from altered amino-acid sequences, we focused on the 522,092 synonymous mutations (**Fig. 7A**).

**Fig. 7.**
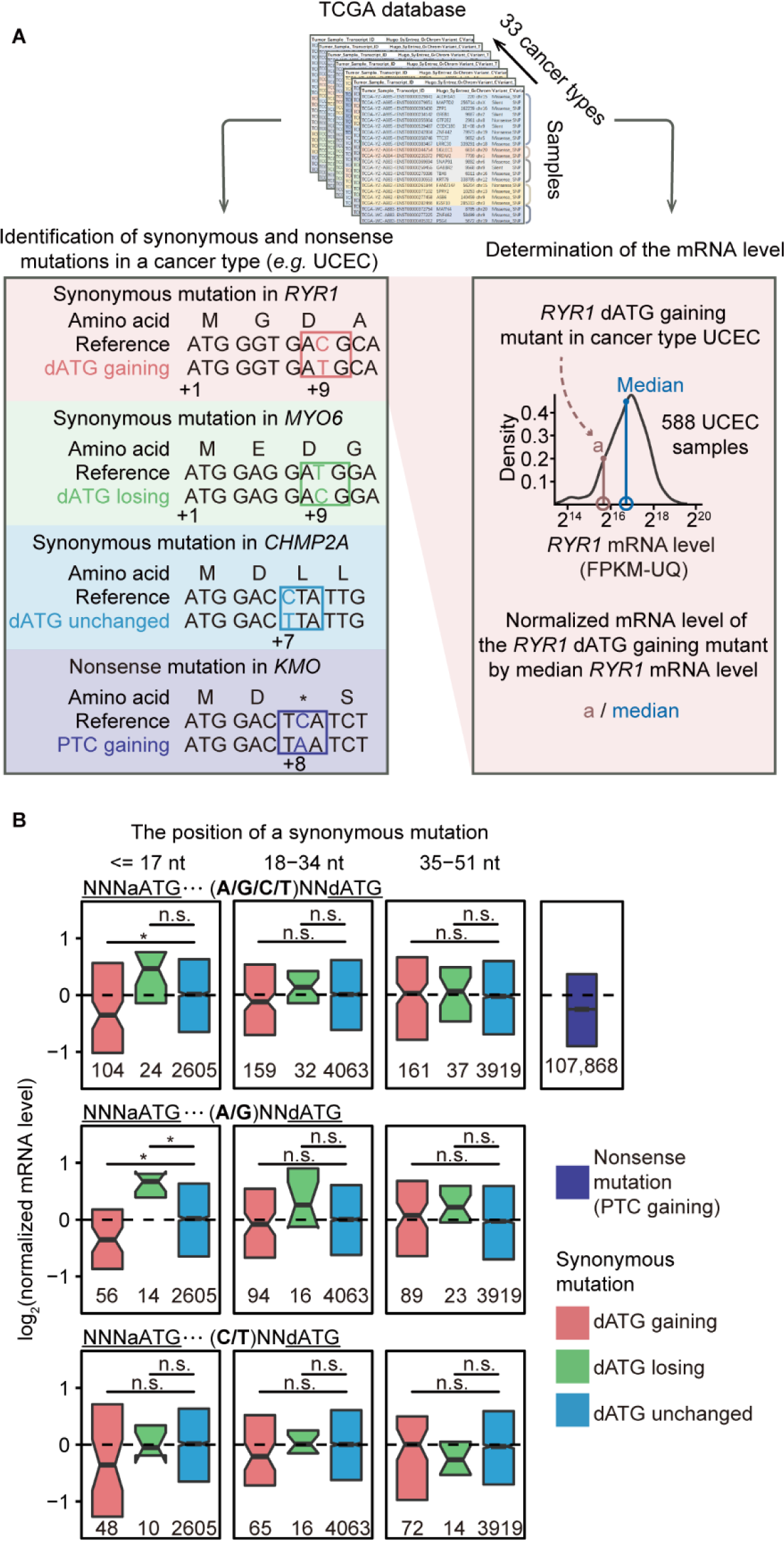
The reduction in the mRNA level caused by somatic synonymous mutations that introduce proximal out-of-frame dATGs. (A) Schematic shows the identification of synonymous mutations that introduce or remove dATGs in the cancer genome and determination of the mRNA level of the mutated gene in the corresponding cancer sample. The synonymous mutations that neither introduce nor remove dATGs (“dATG unchanged”) and the nonsense mutations that introduced premature termination codons (PTC) were also identified. The mRNA level of a mutated gene in a cancer sample was normalized by the median of all other samples of the same cancer type. (B) Boxplots show the mRNA levels of genes bearing synonymous mutations in the cancer samples, grouped by the genomic region where the synonymous mutation occurred (relative to the aATG). Nonsense mutations are also shown (as the positive control); the effect size of reduction in the mRNA level appeared small likely because of the low allele frequency of somatic mutations within a cancer sample. Outliers and whiskers are not shown. *P* values were given by the Mann-Whitney *U* tests. **, *P* < 0.01; *, *P* < 0.05.

To determine the impacts of synonymous mutations that introduce dATGs, we estimated the ratio of mRNA levels of genes bearing synonymous dATG-gaining mutation to median mRNA levels of that gene in all other samples of the same cancer type (**Fig. 7A**). If a synonymous mutation has no effect on the mRNA level, we expect this expression ratio will be 1, while somatic mutations that increase mRNA levels will result in a ratio >1, and somatic mutations that decrease mRNA levels will result in a ratio <1. As a positive control, we identified 107,868 somatic mutations that introduced premature termination codons in this dataset, and determined that this classical scenario for NMD activation reduced the mRNA levels by 16% (*P* < 2.2 × 10^−16^, Mann-Whitney *U* test, **Fig. 7B**).

We then grouped the mutations that introduced out-of-frame dATG in cancer samples according to their positions relative to the aATG: one group included proximal dATGs within 17-nt of the aATG (“proximal”), while two additional groups included dATGs in further downstream regions (“distal”). We found that the gain of proximal out-of-frame dATGs significantly reduced mRNA levels, compared to synonymous mutations that neither introduced nor removed out-of-frame dATGs (*P* = 0.01, Mann-Whitney *U* test), while the distal out-of-frame dATGs did not (**Fig. 7B**). These proximal dATG-gaining mutations reduced mRNA levels by 22%, qualitatively comparable to the reduction in mRNA levels incurred by somatic gains of premature termination codons (16%, **Fig. 7B**). As with dATGs in yeast, the reduction in the mRNA levels was undetectable in cancer cells with somatic mutations that introduced proximal, out-of-frame dATGs in a weak context (C- or T-context, *P* = 0.24, Mann-Whitney *U* test, **Fig. 7B**), indicating that the inhibitory effects on mRNA levels by proximal out-of-frame dAUGs were translation dependent.

We also identified synonymous mutations that removed out-of-frame dATGs in cancer samples (**Fig. 7A**). These proximal dATG-losing mutations elevated mRNA levels by 38%, despite not being statistically significant, likely due to the relatively small sample size (**Fig. 7B**). Nevertheless, the proximal dATG-losing mutations within 17-nt of the aATG elevated mRNA levels when the dATG resided in A- or G-context (**Fig. 7B**). Collectively, these observations suggested that somatic dATG mutations could modulate gene expression levels in cancer cells.

## Discussion

In this study, we quantified the effects of dATG on GFP translation in thousands of yeast variants, which provided us an opportunity to dissect the many determinants of translation initiation (*e.g.*, codon usage and mRNA secondary structure in the proximal sequences downstream of the aAUG) because the effect of confounding factors would be canceled out among variants. We found that proximal out-of-frame dAUG inhibited translation initiation at the aAUG, data that are compatible with a Brownian ratchet scanning process: small-amplitude 5′–3′ and 3′–5′ oscillations result in net 5′–3′ PIC progression. Note that the term dATG used here denotes an ATG downstream of the aATG, different from the ATGs downstream of stop codons as termed in a previous study (Wu et al., 2020).

Statistical analyses revealed a 77% average leakage rate for each single scan of an AUG triplet by the PIC and that the net leakage rate of 8% was achieved by multiple scans of the same triplet. A net leakage rate of 8% may ostensibly seem high considering the volume and variety of functional proteins that require accurate translation to maintain cellular function. However, based on our results, we propose that this leakage rate is reasonable for two reasons. First, the addition of a second in-frame initiation codon as an auxiliary initiation site for translation initiation significantly elevates GFP levels (orange box in **Fig. 1D**), indicating that a substantial fraction of PICs appear to miss the aAUG and are instead captured by an in-frame, downstream AUG (Benitez-Cantos et al., 2020). Second, the canonical ORFs are translated in genes bearing out-of-frame uATGs (Ingolia et al., 2009; Johnstone et al., 2016; Juntawong et al., 2014; Stumpf et al., 2013; Wang and Rothnagel, 2004; Yang et al., 2017; Zhang et al., 2018), which indicates that a substantial proportion of uAUGs are not recognized by PICs.

Our calculation of a 77% average leakage rate for each single scan of an AUG triplet is indeed unexpectedly high. However, this high rate is intuitively consistent with data showing that a dAUG placed at +6 position (with a G at its −3 position) can reduce GFP intensity by ∼40% (*i.e.*, ∼40% PICs that could have initiated at the aAUG were retained by a dAUG 5-nt apart, **Fig. 1E**). Furthermore, when the aATG is in a weak context (with a T at its −3 position), the addition of a dATG at the +6 position even causes a ∼75% reduction in GFP intensity (**Fig. 2A**). Consistently, it was also reported that the proximal, downstream, overlapping ORF in a bicistronic viral mRNA could cause a ∼60% reduction in translation initiation at the aATG, suggesting a substantial single-scan leakage rate (Matsuda and Dreher, 2006).

Previous studies have shown that PICs scan at a rate of ∼8–10 nt/s based on the relationship between the length of 5′-untranslated region (5′-UTR) and the time required for the first round of translation products (Berthelot et al., 2004; Vassilenko et al., 2011). However, it is puzzling that this scanning rate is even lower than the rate of translation elongation, which is ∼6 codons/s, or ∼18 nt/s (Bostrom et al., 1986; Ingolia et al., 2011).

This incongruity is further exacerbated if we consider that PIC scanning only inspects for complementarity to the Met-tRNAi anticodon while translation elongation includes several complicated steps such as tRNA selection, peptide bond formation, and translocation (Schuller and Green, 2018; Zhao et al., 2020). We thus propose that PIC scanning rates measured in previous studies represent the “net” scanning rate, and can be better understood in the framework of the Brownian ratchet scanning model: that is, if all back-and-forth movements along the mRNA by the PIC are taken into account, the actual scanning rate is ∼10 fold faster than the overall net scanning rate, or ∼80–100 nt/s. This scanning rate is four to six times faster than the rate of translation elongation and approximately two times faster than the transcription (∼60 nt/s) and replication (∼50 nt/s) rates (Fuchs et al., 2014; Sekedat et al., 2010).

Some seminal experiments showed that immediately post-termination, ribosomes can scan in the 3′–5′ direction and thereby reinitiate translation at a nearby AUG triplet upstream of the stop codon (Peabody et al., 1986; Skabkin et al., 2013). This finding suggests that the 40S ribosomal subunit has an intrinsic capability to migrate in both the 5′–3′ and 3′–5′ directions along unstructured mRNAs (Kozak, 1979; Pestova and Kolupaeva, 2002; Skabkin et al., 2013). However, these observations differ from our findings reported here, since it remains unknown if the 3–5′ movement described in previous studies only occurs before the post-termination ribosomes have recruited sufficient eIFs required for the “normal” scanning process that starts at the 5′-cap (Kozak, 2001; Skabkin et al., 2013).

While we suggest that the inhibitory effects of dAUG are evidence supporting a 3′–5′ PIC movement, it is possible in principle that this inhibition could be explained, at least in part, by steric effects of ribosomes occupying at dAUGs waiting for translation initiation. That is, a ribosome occupying at nearby downstream location could lead to steric hindrance of aAUG recognition by the PIC (Zhao et al., 2021), thereby reducing the translation initiation rate at the aAUG. However, we propose that our observations are better explained by the Brownian ratchet scanning model, because the inhibitory effect of dAUG on GFP intensity gradually decrease in response to increasing aATG-dATG distance, instead of a stepwise transition as expected in the steric hindrance model (**Fig. 1E** and **Fig. S4B**).

In addition to the strictly unidirectional scanning and Brownian ratchet scanning models that we compared in this study, other researchers have proposed that the PIC can move to the AUG codon via PIC “diffusion” along the mRNA (Berthelot et al., 2004; Vassilenko et al., 2011). The reason why we did not explicitly test the diffusion model here is that prior studies have rejected this hypothesis based on observations that 5′-UTR length and time required for the first round of translation products share a linear relationship, rather than the square relationship as predicted by the diffusion model (Berthelot et al., 2004; Vassilenko et al., 2011). Nevertheless, the diffusion model is actually nested within our Brownian ratchet scanning model, and therefore, can be tested by determining if the probability of adding a pawl (*p.Pawl*) is equal to zero. The parameter *p.Pawl* was estimated to be significantly >0 (0.1% with the standard error equal to 0.02%, **Fig. 5F**) in our MCMC analyses, again supporting a Brownian ratchet model for PIC scanning rather than a diffusion model.

Furthermore, it remains controversial as to whether the 5′-cap remains attached to the scanning PIC after recruitment of the small ribosomal subunit (Jackson et al., 2010; Shirokikh and Preiss, 2018). A cap-severed scanning model proposes that multiple PICs can scan a 5′-UTR at the same time (Berthelot et al., 2004), whereas a cap-tethered scanning model hypothesizes that only one PIC at a time can scan the 5′-UTR (Bohlen et al., 2020). The Brownian ratchet scanning model is intrinsically more compatible with the cap-tethered scanning model because the pawls placed on the 5′-UTR must dissociate from mRNA at a rate that is sufficiently slow to restrict 3′–5′ PIC movement beyond the pawl, which in turn, could block the 5′–3′ directional scanning by a following PIC (if present). Furthermore, under the cap-tethered scanning model there would be only one PIC scanning for AUG at the same time, ruling out the possibility of interference among multiple oscillating PICs along the same mRNA.

Our findings in this study suggest at least three major directions for future experimental exploration. First, it would be of great value to confirm or refute the Brownian ratchet model by tracing the movement of a single PIC along an mRNA with super-resolution light microscopy (Betzig and Chichester, 1993) or optical tweezers (Ashkin et al., 1986), in real-time and at single-nucleotide resolution, to try to observe small-amplitude 5′–3′ and 3′–5′ oscillations with a net 5′–3′ movement. Second, the quantification of the consumption of adenosine triphosphate (ATP) usage for PIC scanning would also be mechanistically informative (Brito Querido et al., 2020; Pestova and Hellen, 1999) since a strictly unidirectional scanning model predicts that the number of consumed ATPs will be an integer multiple of 5′-UTR length because ATPs should be used in each movement (one nt per step) to ensure directionality; by contrast, the Brownian ratchet scanning model predicts that fewer ATPs will be consumed than 5′-UTR length because ATPs are only used occasionally for pawl placement onto mRNAs (Spirin, 2009). Third, the identification of eukaryotic initiation factors that participate in the Brownian ratchet scanning mechanism (*e.g.*, the protein identity of the “pawl”) could also offer insight into the mechanism by which directional PIC movement can be achieved from PIC diffusion in both 5′–3′ and 3′–5′ directions.

In summary, our results provide quantitative support for the Brownian ratchet model of PIC scanning, and show how such a process can influence genome evolution. In addition, our findings also have medical implications. Specifically, the currently common presumption by experimental biologists and medical scientists that disease-associated, translational defect mutations are likely associated with uAUGs (Calvo et al., 2009; Zhang et al., 2019) should be expanded to include inhibitory effects of proximal, out-of-frame dAUGs on the translation of canonical ORFs. This new insight will help in computational predictions of disease-causing mutations from whole genome/exome sequencing data in the future.

## STAR Methods

### KEY RESOURCE TABLE

| REAGENT or RESOURCE | SOURCE | IDENTIFIER |
| --- | --- | --- |
| Biological samples |  |  |
| *S. cerevisiae* |  | N/A |
| Chemicals, peptides, and recombinant proteins |  |  |
| G418 | Amresco | Cat#97063-058 |
| Nourseothricin | Amresco | Cat#6021-878 |
| Hygromycin B | Amresco | Cat#97064-454 |
| Glass beads, acid-washed | Sigma-Aldrich | Cat#G8772 |
| Yeast Extract | Thermo Scientific | Cat#LP0021 |
| Bacto peptone | BD | Cat#211677 |
| D(+) galactose | VWR | Cat#97062-080 |
| Yeast uracil minus media | FunGenome | Cat#YGM003A-3 |
| Dextrose (glucose) | SCR | Cat#10010518 |
| Glycerol | SCR | Cat#10010618 |
| Ethanol | SCR | Cat#10009218 |
| Critical commercial assays |  |  |
| Phusion® High-Fidelity DNA Polymerase | NEB | M0530 |
| GeneArt seamless cloning and assembly enzyme mix | Invitrogen | Cat#A14606 |
| GoScript™ Reverse Transcription System | Promega | Cat#A5001 |
| Dual-Luciferase® Reporter Assay System | Promega | Cat#E1910 |
| Deposited data |  |  |
| FACS-seq data | This paper | CRA005203 |
| RNA-seq data | This paper | CRA005203 |
| DNA-seq data | This paper | CRA005203 |
| SSU data | Archer et al. (2016) | PMID: 27437580 |
| The Cancer Genome Atlas | TCGA |  |
| *Saccharomyces cerevisiae* coding sequences | Ensembl | R64-1-1 |
| *Homo sapiens* coding sequences | Ensembl | GRCh38 |
| *Escherichia coli* coding sequences | GenBank | NZ_CP027599.1 |
| *Bacillus Subtilis* coding sequences | GenBank | NC_000964.3 |
| uATG data | Cuperus et al. (2017) | PMID: 29097404 |
| Experimental models: Cell lines |  |  |
| Human:HeLa cell | ATCC | CCL-2 |
| Experimental models: Organisms/strains |  |  |
| *S. cerevisiae*: Strain background: BY4742 | Brachmann et al. (1998) | PMID:9483801 |
| Oligonucleotides |  |  |
| Primers see Table S1 and Table S2 | This paper | N/A |
| Recombinant DNA |  |  |
| pAG25 | Goldstein and McCusker (1999) | RRID: Addgene_35121 |
| pFA6a-GFP(S65T)-kanMX6 | Bahler et al. (1998) | RRID: Addgene_39292 |
| pUC57 | Markausakas et al. (unpublished) | GenBank: Y14837.1 |
| pmirGLO | Promega | Cat#E1330 |
| Software and algorithms |  |  |
| R | R Development Core Team (2016) | https://www.r-project.org/ |
| RNAfold | Lorenz et al. (2011) | http://rna.tbi.univie.ac.at/ |

### RESOURCE AVAILABILITY

#### Lead contact

Further information and requests for resources and reagents should be directed to and will be fulfilled by the lead contact, Wenfeng Qian

#### Materials availability

All reagents generated in this study are available from the lead contact.

#### Data and code availability

FACS-seq, RNA-seq, and DNA-seq data have been deposited at Genome Sequence Archive (Wang et al., 2017) under the accession number CRA005203 and are publicly available as of the date of publication. Any additional information required to reanalyze the data reported in this paper is available from the lead contact upon request.

### EXPERIMENTAL MODEL AND SUBJECT DETAILS

The *Saccharomyces cerevisiae* strain BY4742 were cultured in 1% yeast extract–2% peptone–2% dextrose (YPD) medium at 30°C. HeLa cells were cultured in Dulbecco’s Modified Eagle Medium containing 10% Fetal Calf Serum, 2 mM L-glutamine at 37°C in a 5% CO_2_ incubator.

### METHOD DETAILS

#### Construction of the yeast dATG library

We constructed the yeast dATG library as described in a previous study (Chen et al., 2017). Specifically, we constructed a yeast strain (BY4742-dTomato, *MAT*α *his3Δ1 leu2Δ0 lys2Δ0 ura3Δ0 gal7Δ0::dTomato-hphMX*) that expressed dTomato from the *GAL7* promoter in the background of BY4742, using a recombination-mediated polymerase chain reaction (PCR)-directed allele replacement method (primers provided in **Table S2**). The dTomato expression would be later used to normalize GFP intensity, which in principle could be affected by cell-to-cell variation in galactose induction and cell-cycle status (Chen et al., 2019). We selected the transformants on the 1% yeast extract–2% peptone–2% dextrose (YPD) solid medium with 200 ug/ml G418 (Amresco). We performed PCR on the extracted genomic DNA of the yeast transformants to verify the successful *GAL7* deletion and *dTomato* integration (here and also hereafter when genetic manipulation was performed).

To construct the dATG yeast library in the background of *Δupf1* (BY4742-dTomato-*Δupf1*), we deleted *UPF1* in the background of BY4742-dTomato using recombination-mediated PCR-directed allele replacement method (primers provided in **Table S2**). To this end, we amplified the *natMX* cassette from plasmid PAG25, transformed the PCR product into BY4742-dTomato, and selected the transformants on the YPD solid medium with 100 ug/ml nourseothricin (Amresco). To control for the potential cellular effect of *natMX* expression when the expression of *GFP* in the background of *Δupf1* was compared with the wild type, we replaced the gene encoding homothallic switching endonuclease, *HO* (a pseudo gene in the BY4742 background), using *natMX* (Wu et al., 2017). The resultant yeast strain (*i.e.*, BY4742-dTomato-*Δho*) was treated as the wild type in this study.

We chemically synthesized 29 oligos with specific positions using doped nucleotides (**Fig. 1B**, **Fig. S1A**, and **Table S1**); 28 of them contained an ATG designed at a particular position downstream of the aATG. We mixed these oligos and fused them with the *GAL1* promoter, the full-length *GFP* coding sequence (CDS, with the initiation codon of *GFP* omitted), the *ADH1* terminator, *URA3MX*, and the *GAL1* terminator (in the order shown in **Fig. S1A**), using fusion PCR and the GeneArt Seamless Cloning and Assembly Kit (Thermo Fisher A14606). The resultant sequence surrounding the synthetic oligo is CTT TAA CGT CAA GGA GAA AAA (NNN NNN ATG NNN NNN NNN NNN NNN NNN NNN NNN NNN NNN) GCA GGT CGA CGG ATC CCC GGG Tta aTt aaC AGt aaA GGA GAA GAA CTT TTC ACT GGA GTT GTC CCA ATT CTT GTt gaA Tta g<u>AT G</u>Gt g<u>aT G</u>Tt a<u>aT G</u>GG CAC AAA TTT, where out-of-frame dATGs and out-of-frame stop codons are underlined and in lower case letters, respectively.

To construct the dATG yeast library, we transformed the PCR product to replace the coding sequence of *GAL1* in the BY4742-dTomato-*Δho* strain (**Fig. S1A**). Specifically, the *GAL1* promoter (500 nt) and *GAL1* terminator (500 nt) were used as long homologous sequences to allow efficient integration of the PCR product into the yeast nuclear genome (Chen et al., 2017). We selected successful transformants in the synthetic complete medium (dextrose as the carbon source) with uracil dropped out because *URA3* was used as the marker gene. We collected a total of ∼50,000 yeast transformants, which most likely contained various sequences surrounding the aATG of *GFP*, due to the huge number of possible sequences that could be generated from the synthesized oligos (4^36^ ≈ 5 × 10^21^). We similarly constructed the dATG yeast library in the background of BY4742-dTomato-*Δupf1*, and collected a total of ∼60,000 yeast transformants.

#### FACS-coupled high-throughput sequencing (FACS-seq)

We gauged GFP intensities for each of the thousands of yeast dATG variants using a high-throughput strategy, FACS-seq, as described in a previous study (Chen et al., 2017). Specifically, we pooled yeast variants that contained various sequences surrounding the aATG of *GFP* and cultured them in the liquid medium (YPGEG) that contained 1% yeast extract, 2% peptone, 2% glycerol, and 2% ethanol (both serve as the carbon source), and 2% galactose (to induce the expression of *GFP* and *dTomato*). We harvested yeast cells after 18 hours, in which duration the optical density at 660 nm increased from ∼0.1 to ∼0.7. We nitrogen froze half of the harvested cells for total RNA and DNA extraction (to perform RNA-seq and DNA-seq as described below), and re-suspended the other in the 1 **×** phosphate buffered saline for FACS-seq.

We sorted yeast cells into eight bins using Aria III cytometer (BD Biosciences) based on the intensity ratio of GFP and dTomato fluorescence, which were excited by 488 nm and 561 nm lasers, and were detected using 530/30 nm and 610/20 nm filters, respectively. We recorded the median GFP/dTomato intensity ratio for each bin, as well as the proportion of cells belonging to the “gate” of each bin (**Fig. S1A**). We collected at least 20,000 yeast cells for each bin and cultured them individually in YPD overnight at 30°C to amplify the cell population. Since GFP was not expressed in YPD (therefore should confer limited fitness cost), the relative fraction of yeast variants in each bin should be largely maintained.

We extracted the genomic DNA from yeast cells of each bin and performed two rounds of PCR amplification on the variable region (*i.e.*, 6-nt upstream and 30-nt downstream of the *GFP* aATG) to construct the Illumina sequencing libraries. Briefly, in the first round PCR, a pair of sequences identical to the 21-nt sequence upstream of aATG (positions −35 to −15, relative to the A[+1] of the aATG) and the reverse complement of the 20-nt sequence downstream of the aATG (positions +45 to +64) were used to amplify the variable region (primers provided in **Table S2**). Meanwhile, 19-nt or 21-nt sequences identical to the 3′-end of the P5 or P7 adaptor, respectively, a 12-nt stretch of random nucleotides (NNNNNNNNNNNN, designed to avoid difficulty in base calling of Illumina sequencing when sequencing “constant” region), and a 6-nt bin specific barcode were introduced to the ends of the PCR product (**Fig. S1A**). In the second round PCR, the full-length P5 and P7 adaptors as well as the sequencing indices were added to the ends. The PCR products were then subject to Illumina sequencing (NovaSeq 6000 platform, in the PE150 mode).

#### Small-scale validation of fluorescence intensities using flow cytometer

We randomly isolated 20 yeast variants from the yeast dATG library from individual colonies on the solid medium, and sequenced the variable region for each variant by Sanger sequencing. We induced the expression of the fluorescent proteins in the liquid YPGEG media for individual strains, and harvested yeast cells in the mid-log phase. For each strain, we measured the GFP and dTomato fluorescence intensities by Aria III cytometer (BD Biosciences), which were excited by 488 nm and 561 nm lasers and detected with 530/30 nm and 610/20 nm fluorescence filters, respectively.

#### RNA-seq and DNA-seq for the yeast dATG library

We extracted the total RNA from the harvested cells of the yeast dATG library (cultured in the YPGEG liquid medium for 18 hours), and performed reverse transcription using the GoScript^TM^ Reverse Transcription System (Promega). We built the Illumina sequencing library by two-round PCR-amplification of the variable region (similarly to FACS-seq, primers provided in **Table S2**). Illumina sequencing was performed on the NovaSeq 6000 platform under the PE150 mode. To control for the variation in the cell number among the yeast dATG variants and the potential bias in Illumina sequencing, we also extracted the total genomic DNA from the harvested cells of the yeast dATG library and PCR-amplified the variable region for Illumina sequencing (primers provided in **Table S2**).

We performed two biological replicates by independently inducing the GFP expression in the YPGEG liquid medium for cells in the yeast dATG library. In each replicate we performed FACS-seq, RNA-seq, and DNA-seq for the same group of harvested cells.

#### Dual-fluorescence reporter assay in yeast

We determined the distance-dependent inhibitory effect of out-of-frame dAUGs on translation initiation at the aAUG in small-scale experiments, using a dual-fluorescence reporter assay. To this end, we first constructed *TEF* promoter-*GFP* CDS-*CYC1* terminator-*TDH3* promoter-*dTomato* CDS-*ADH1* terminator-*URA3MX* in the background of pUC57 plasmid using the GeneArt Seamless Cloning and Assembly Kit (Invitrogen, A14606, **Table S3**). Then, with this plasmid as the template, we introduced an out-of-frame dATG at the position +8, +14, +20, or +26 by inserting a 27-nt sequences downstream of the aATG of *GFP* using fusion PCR (primer sequences provided in **Table S2**). Note that the four versions of the 27-nt sequences encode the same nine amino acids (*i.e.*, the dATGs were generated from synonymous mutation, **Fig. 1F**).

We inserted each of the four dATG variants into the BY4742 genome by replacing the endogenous *HO* locus in Chromosome IV, using recombination-mediated PCR-directed allele replacement method (59-nt homologous sequences in both ends, primers are provided in **Table S2**). We harvested yeast cells in the mid-log phase and used Accuri^TM^ C6 cytometer to measure the GFP and dTomato fluorescence (excited by 473 nm and 552 nm lasers and detected with 530/30 nm and 610/20 nm filters, respectively). The reported GFP fluorescence intensity was normalized by the dTomato fluorescence intensity.

#### Overlapping dual-fluorescence reporter assay in yeast

To detect the translational competition between two closely spaced AUGs we designed an overlapping dual-fluorescence reporter, in which GFP (frame 0) and dTomato (frame +1) were encoded in the same transcript, which were expressed from the *TDH3* promoter, (**Fig. 2C**). To avoid truncated protein in frame +1, we removed all stop codons in frame +1 in the *GFP* CDS via synonymous mutations. To minimize the influence of the long peptide in the N-terminus (peptide sequence encoded by frame +1 of the *GFP* CDS) on protein folding of dTomato, we inserted a 2A self-cleaving peptide (Souza-Moreira et al., 2018) in frame +1 right upstream to the *dTomato* CDS. The overlapping dual-fluorescence reporter DNA was synthesized by BGI Tech (sequence shown in **Table S3**).

We inserted the overlapping dual-fluorescence reporter into the genome, which replaced the endogenous *HO* locus in BY4742, by recombination-mediated PCR-directed allele replacement method (59-nt homologous sequences in both ends, primers provided in **Table S2**). We constructed two reporters with 3-nt difference in the sequence upstream of the dTomato ATG (**Fig. 2C**). We harvested yeast cells in the mid-log phase and measured the GFP and dTomato fluorescence using Accuri^TM^ C6 cytometer.

#### Dual-luciferase assay in HeLa cells

To detect the inhibitory effect of proximal out-of-frame dAUGs on translation initiation at the aAUG in HeLa cells, we performed a dual-luciferase assay based on modified pmirGLO plasmids, in which firefly and *Renilla* luciferases were individually expressed from *PGK* and SV40 promoters, respectively. Specifically, using site-directed mutagenesis methods, we modified the pmirGLO plasmids (from Addgene) by introducing a 6-nt sequence (AATTTT, weak context) right upstream of the firefly luciferase ATG to increase its leakage rate. We designed synonymous mutations to generate four 21-nt sequences that encode the same amino acid sequence; one sequence did not contain proximal out-of-frame dATGs and the other three each contained a proximal out-of-frame dATG at the +8, +14, or +20 position relative to the aATG (**Fig. 6F**).

We introduced each of the four 21-nt sequences right downstream of the firefly luciferase ATG in the plasmid using site-directed mutagenesis methods, and individually transfected each of the four modified plasmids into HeLa cells using Lipofectamine 2000 (Thermo Fisher). We determined the activities of luciferases in 96-well microliter plates 48 hours after transfection, using a commercial dual luciferase assay kit (Promega, E1910) following the manufactory’s protocol. Briefly, we lysed HeLa cells using 500 μL of passive lysis buffer and mixed 20 μL suspension with 100 μL firefly luciferase substrate. We first measured firefly luciferase activity using the Synergy HTX multi-mode microplate reader (BioTek). Then, we added 100 μL of Stop-and-Glo reagent to the solution, and measured the *Renilla* luciferase activity using the same equipment.

### QUANTIFICATION AND STATISTICAL ANALYSIS

#### Quantification of GFP and mRNA levels for each dATG variant

The “read 1” of a read pair from the DNA-seq data should follow the pattern of N(12)-barcode (6 nt)-CCTCTATACTTTAACGTCAAGGAGAAAAA-N(6)-ATG-N(30)-GCAGGTCGACGGATCCCCGGGTTAATTAACA-barcode (6 nt)-N(12)-P7. Note that P7 adaptor would also be sequenced downstream to the full length of the inserted sequence because the length of the inserted sequence of the Illumina sequencing library generated in this study was 135 nt, shorter than the length of a sequencing read (150 nt). For the same reason, the reverse complements of the barcodes and the variable region would be sequenced for a second time in the “read 2”, in which part of the P5 adaptor would be sequenced. For each read, we extracted the barcodes (6-nt upstream + 6-nt downstream of the variable region) as well as the 36-nt variable sequence surrounding the ATG using pattern matching. We discarded the whole read pair in the following three cases: 1) if either read of a read pair could not match to the pattern, 2) if any of the four barcodes extracted from a read pair was different from the barcodes that were introduced during library preparation for a particular sample (**Table S2**), or 3) if the read 1 sequence and the reverse complement sequence of the read 2 were not identical in the variable region. We then classified read pairs into biological replicates according to the barcode sequence, and grouped read pairs into variants according to the sequence in the variable region. The sequencing data from the RNA-seq and FACS-seq libraries were similarly analyzed. The numbers of read pairs that passed the three criteria and the numbers of identified variants are summarized in **Table S4**.

Some sequences in the variable regions were not detected in all three libraries (FACS-seq, RNA-seq, and DNA-seq), which implied that they were potentially originated from PCR-amplification errors during Illumina sequencing library preparation. We therefore discarded the dATG variants that did not appear in all three libraries. Furthermore, the frequencies of some dATG variants appeared to be too low in the DNA-seq (number of read pairs ≤8) and FACS-seq libraries (all read pairs from the eight bins combined ≤64). To be conservative, we also discarded these variants (the remained variant numbers are shown in the Venn diagram of **Fig. S1D**). Additional filtering criteria are listed in **Fig. S1D**. In particular, variants containing in-frame stop codons in the 30-nt downstream regions showed lower GFP intensities (**Fig. S1E**) as they are potential NMD substrates; we removed these variants from the subsequent analyses. Furthermore, we also discarded the variants containing uATG due to their potential impact on translation initiation of GFP (**Fig. S1F**).

Following a previous study (Chen et al., 2017), the dTomato-normalized GFP intensity of each yeast variant (*Gj*) was calculated as the average GFP/dTomato intensity ratio among the eight bins, weighted by the proportions of its cells distributed in the eight bins (**Fig. S1A** and **Table S5**). The weight of variant *j* in bin *i* was estimated from *P_i_ × n_ij_*, where *n_ij_* was the fraction of read pairs for variant *j* among all read pairs in bin *i*, and *P_i_* was the proportion of cells belonging to the “gate” of bin *i* as recorded by the flow cytometer.

The GFP level of variant *j* was calculated from the formula:

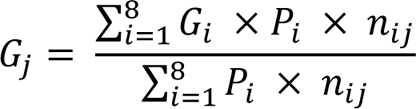

where *G_i_* was the median GFP/dTomato ratio estimated from the collected yeast cells in bin *i* by the flow cytometer. We obtained GFP intensities for a total of 35,582 variants, and the GFP intensity of each variant is provided in **Tables S6**.

We estimated the mRNA level for each dATG variant from its read pair frequencies in the RNA-seq and DNA-seq libraries. Specifically, the read pair frequency of variant *i* in the RNA library (*R_i_*) or DNA-seq library (*D_i_*) was calculated from the fraction of read pairs derived from variant *i* among the total number of read pairs in the RNA-seq or DNA-seq library, respectively. Then, the mRNA level (abundance per cell) of variant *i* was estimated from the *R_i_*/*D_i_* ratio. The mRNA level of each variant is provided in **Tables S6** and **Tables S7**.

#### Estimation of leakage rates from the small ribosomal subunit (SSU) footprint data

SSU footprint data in yeast was retrieved from a previous study (Archer et al., 2016). The SSU reads whose 5′-ends were located in the 40-nt upstream (uSSU) and 40-nt downstream (dSSU) region of the aATG for each gene were counted. The leakage rate of a gene was estimated from the fraction of downstream footprints among all footprints detected within the ± 40-nt region (*i.e.*, # of dSSU / [# of uSSU + # of dSSU]). The leakage rates were also estimated for the SSU footprints within the ± 80-nt region (**Fig. S2C**).

#### Estimation of minimum free energy (MFE) and codon adaption index (CAI)

The mRNA secondary structure right downstream of ATG was reported to regulate translation initiation/elongation (Kudla et al., 2009; Yang et al., 2014). Since the intrinsic propensity of RNA sequences to form a secondary structure could be inferred from MFE, we estimated MFE in the 30-nt region downstream of the aATG for each variant in our dATG library using the RNAfold command (RNAfold -d2 --noLP --noPS) in the package ViennaRNA (Lorenz et al., 2011).

The synonymous codon usage was also reported to regulate protein synthesis rate (Chu et al., 2014). We therefore calculated CAI in the 30-nt (10-codon) region downstream of the aATG for each dATG variant, following the computational procedure described in previous studies (Sharp and Li, 1987; Yang et al., 2017).

#### Assessment of distance-dependent translational inhibition by uATGs

The growth rate data (in the medium without histidine) for yeast variants that harbor uATGs upstream of *HIS3* were retrieved from a previous study (Cuperus et al., 2017), from which the protein abundance of HIS3 was inferred for each variant. Only the yeast variants that harbor one uATG (located in the 3–29-nt upstream region of aATG) were used in the subsequent analyses. Yeast variants that bore uATGs located in the 3–5-nt upstream region of aATG were discarded because the presence of these uATGs constrained the sequence context (nucleotide at the −3 position) of the aATG.

#### Simulation of Brownian ratchet scanning process using a random walk model

We simulated the ratchet-and-pawl mechanism using a modified random walk model (Codling et al., 2008). The PIC occupies 19 nt in the mRNA, and among them, the triplet at the 13^th^–15^th^ position is inspected for complementarity to the Met-tRNAi anticodon (Archer et al., 2016). The PIC starts out with its 5′-trailing side at the 5′-cap and takes one nt per step along the mRNA. A pawl is stochastically placed onto the mRNA at the 5′-trailing side of the PIC with the probability of *p.Pawl*, and the disassociation of the pawl from the mRNA is sufficiently slow that is not considered in the model. The PIC moves with equal probabilities in the 5′–3′ and 3′–5′ directions (each 50%) unless its 5′-trailing side hit the 5′-cap or a pawl, in which circumstances, the PIC moves in the 5′–3′ direction with 100% probability. When an AUG is located within the 13^th^–15^th^ positions covered by the PIC, the PIC may recognize the AUG in the probability of (1 − *p.Leakage*) or miss the AUG in the probability of *p.Leakage*; in the latter case, the PIC continues scanning. Note that in our simulation we assume that AUG triplets can be recognized when the PIC moves in either the 5′–3′ or 3′–5′ direction. Sometimes the PIC may recognize out-of-frame AUGs, which would activate the NMD pathway and mRNA degradation with the probability of *p.NMD*.

We simulated the PIC scanning processes on one Solo and 25 Duo variants. The Solo variant contained 50-nt sequence upstream and 50-nt sequence downstream of the aATG. The distance between the two ATGs in one of the 25 Duo variants was within the range of 6–30 nt, and the Duo variants also contained 50-nt sequence upstream of the aATG and 50-nt sequence downstream of the dATG. When the PIC moved beyond the 50-nt downstream region (*i.e.*, beyond the 3′-end), NMD was also activated with the probability of *p.NMD* as the PIC might encounter out-of-frame AUGs further downstream (the next three ATG triplets downstream of the variable region of *GFP* are all out-of-frame). We simulated the PIC scanning process for each variant (the Solo or one of the 25 Duos) 100 times, and calculated the fraction of successful translation initiation events at the aATG. The protein expression level of a variant was estimated from the product of this fraction and the proportion of mRNA that did not activate the NMD pathway. We then compared this protein expression level in our simulation (**Fig. 5D**) with the GFP intensities measured in the experiments (variants of the two replicates combined), by calculating the residual sum of squares (RSS). Note that translation initiation at the in-frame dAUGs in Duo variants would synthesize functional proteins and would not activate NMD (**Fig. 5A**).

#### Estimation of *p.Pawl, p.Leakage,* and *p.NMD* by the MCMC algorithms

To determine the parameters to start with in the MCMC algorithms, we screened the parameter space for *p.Pawl*, *p.Leakage*, and *p.NMD*. Specifically, we set each of the probability parameters as one of the numbers in 0.001, 0.01, 0.1, 0.2, 0.3, 0.4, 0.5, 0.6, 0.7, and 0.8 (**Fig. S6A**). Together there were (10^3^ =) 1000 combinational value sets. Each parameter set was individually used to simulate the Brownian ratchet scanning process, and then, the RSS was estimated from the simulated protein expression levels and observed GFP intensities in the yeast experiments. Note that the protein expression levels of Duo variants were normalized by that of the Solo variant in both simulation and experimentation, so that the protein levels were directly comparable.

To estimate *p.Pawl*, *p.Leakage*, and *p.NMD*, three hundred iterations of the MCMC simulation were performed. In each iteration, one of the three parameters was changed sequentially in the order of *p.Pawl*, *p.Leakage*, and *p.NMD*. We set the sampling window width for each parameter as 0.0004, 0.2, and 0.5, respectively, which were proximately two times the standard deviation in the 10 parameter sets obtained in the initiation screening (**Fig. S6B**). We generated a new parameter randomly based on the uniform distribution in the window defined by the sampling window width, centered at the parameter value in the previous iteration. If the boundary of the window was smaller than 0 or greater than 1, the boundary was set as 0 or 1 since all three parameters were probabilities.

The new parameter set was used to simulate the PIC scanning processes. If the RSS of the current iteration was smaller than the previous iteration, the previous set of parameters was replaced by the new set of parameters, whereas if the RSS increased, the previous set of parameters remained. After 300 iterations, the parameter set that exhibited the minimum RSS values was recorded. We repeated the simulation 30 times, and obtained the distribution of the three parameters as shown in **Fig. 5E** and **Fig. S6C**. The average value and the standard errors (SE) were estimated from the 30 sets of optimized parameters by the MCMC algorithms (**Fig. 5F**). The net leakage rate (*i.e.*, the fraction of PICs that eventually miss an AUG after multiple scans) was estimated from the fraction of PICs that failed to initiate translation at the aATG in the Solo variant.

To estimate the *p.Leakage* for the ATG in the A-, G-, C-, and T-contexts individually, we performed the MCMC algorithms with 100 iterations, during which we estimated the RSS between the simulated protein expression levels and the GFP intensities measured for the Solo variants that with A, G, C, or T at the −3 position in the experiments. In the simulation, we fixed the *p.Pawl* and *p.NMD* values as acquired above (*p.Pawl* = 0.001 and *p.NMD* = 0.62). The MCMC algorithms were performed 30 times, and average *p.Leakage* for each of the four ATG contexts was estimated from the average value of the 30 MCMC chains (**Fig. 5G**).

#### Detection of out-of-frame dAUGs in eukaryotic and prokaryotic genomes

The coding sequences of the budding yeast (*Saccharomyces cerevisiae*, R64-1-1) and humans (*Homo sapiens*, GRCh38) genes were downloaded from the Ensembl database (Yates et al., 2020) using BioMart (Kinsella et al., 2011), and the sequence of the main transcript of each gene defined in the Ensembl database were used for the subsequent analyses. A gene was discarded if its annotated initiation codon was not ATG or if it was shorter than 45 nt in length. The numbers of genes used for the subsequent analyses were 6564 (yeast) and 19,496 (humans). The genes harboring dATGs located at each position within the 45-nt region downstream of the aATG were counted in each genome.

As a negative control, we also retrieved the coding sequences of two prokaryotic genomes (NCBI Resource Coordinators, 2018), *Escherichia coli* (GenBank: NZ_CP027599.1) and *Bacillus Subtilis* (GenBank: NC_000964.3), where no scanning mechanism is required for recognition of the initiation codon. We applied the same set of criteria as described above for yeast and humans (except for the choice of the main transcript due to the absence of introns) and identified 5113 (*E. coli*) and 3282 (*B. Subtilis)* genes for the subsequent analyses.

#### Determination of the impact on the mRNA level of dATG-changing somatic mutations in cancer samples

Somatic mutation data for 33 cancer types were downloaded from TCGA using the *R* package TCGAbiolinks (Mounir et al., 2019), which were detected by the MuTect2 pipeline (Cibulskis et al., 2013). Somatic mutations that introduced/removed out-of-frame dATGs were identified based on human reference genome GRCh38, and only synonymous mutations was used for the subsequent analyses, in an effort to exclude the potential interference from amino-acid sequence alterations caused by somatic mutations. The nucleotide at the −3 position of the aATG of each human gene was also extracted from the reference genome.

The RNA-seq data were also obtained from TCGA in the unit of FPKM-UQ (Per Kilobase of transcript per Million mapped reads upper quartile). The mRNA level of genes bearing synonymous mutation was normalized by the median mRNA levels of that gene in all other samples of the same cancer type.

#### Supplemental information

**Fig. S1.** High-throughput quantification of GFP intensities by FACS-seq, related to Fig.1and **Fig. S2.** Frame- and context-dependent inhibitory effects on protein synthesis by proximal dAUGs, related to Fig. 1 and Fig. 2. **Fig. S3.** Codon adaption index (CAI) and minimum free energy (MFE) are not significantly varied among Duo variants with dATGs at various locations, related to Fig.2. **Fig. S4.** Two mechanistic explanations of the inhibitory effects on protein synthesis by proximal dAUGs, related to Fig. 2. **Fig. S5.** High-throughput measurement of mRNA levels for dATG variants, related to Fig. 3. **Fig. S6.** The optimization of the parameters in the Brownian ratchet scanning model using the MCMC algorithms, related to Fig. 5. **Fig. S7.** The comparison of the observed GFP intensities of Duo variants in the yeast experiments and the simulated GFP intensities for each of the 30 sets of optimized parameters by the MCMC algorithms, related to Fig. 5.

**Table S1.** Doped nucleotide oligos used to construct the yeast dATG library, related to Fig. 1, Fig. S1, and STAR Methods. **Table S2.** Primers used in this study, related to STAR Methods. **Table S3.** Coding sequences for the dual-fluorescence reporter and the overlapping dual-fluorescence reporter, related to Figs. 1F, Figs. 2C, and STAR Methods. **Table S4.** Number of variants in each sample, related to STAR Methods. **Table S5.** Information for individual bins in FACS-seq, related to Fig. 1, Fig. S1, and STAR Methods. **Table S6.** Sequence, GFP intensity, and mRNA level of the yeast variants generated in the background of *Δho*, related to STAR Methods. **Table S7.** Sequence and mRNA level of yeast variants in the background of *Δupf1*, related to STAR Methods.

## Acknowledgments

We thank Dr. Xiaolei Su and Dr. Junjie Guo from Yale University, Dr. Lucas Carey from Ginkgo Bioworks, Dr. Mengyi Sun from Northwestern University, Dr. Yuanchao Xue from Institute of Biophysics CAS, and Dr. Yuqiang Jiang, Dr. Zhuo Du, and Dr. Qiang Tu from Institute of Genetics and Developmental Biology CAS for discussion. This work was supported by grants from the National Key Research and Development Program of China (2019YFA0508700) and the National Natural Science Foundation of China (31922014 and 32100443).

## Authors’ contributions

W.Q. designed the study; K.L., S.Z., and T.Z. performed experiments; J.K. and S.Z. performed computational analyses; K.L., J.K., S.Z., and W.Q. wrote the manuscript.

