## Supplemental information for "Distance-dependent inhibition of translation initiation by downstream out-of-frame AUGs reveals that ribosome scans in a Brownian ratchet process"

Supplemental information includes Supplemental Figures S1–7 and Supplemental Tables S1–7.

### Supplemental Figures

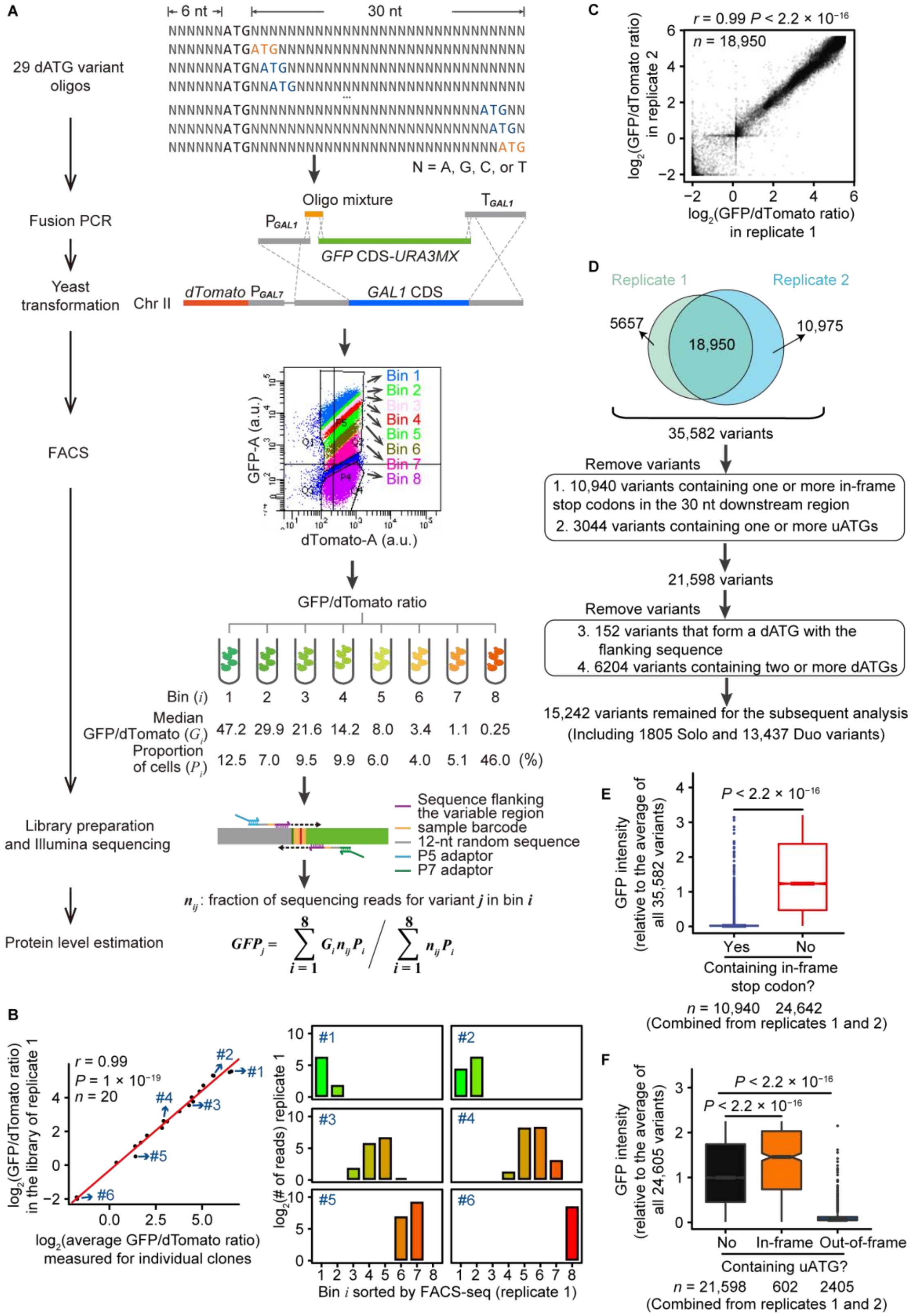


$$GFP_j = \sum_{i=1}^{8} G_i n_{ij} P_i \Big/ \sum_{i=1}^{8} n_{ij} P_i$$

**Fig. S1. High-throughput quantification of GFP intensities by FACS-seq, related to Fig. 1.**

(A) Schematic shows the experimental procedure for estimating GFP intensity. The scatter plot shows the GFP and dTomato intensities (area) of individual yeast cells (from biological replicate 1) and the “gates” for sorting yeast cells into eight bins by FACS.

(B) Comparison of the GFP intensities of 20 randomly chosen yeast variants measured by FACS-seq vs. measured individually by flow cytometer on isolated clones (i.e., the average GFP/dTomato ratio among >19,000 cells for each clone). The standard major axis is shown in the red line. Pearson’s correlation coefficient r and the corresponding *P* value are also shown. The six histograms on the right show the number of reads detected in the high-throughput sequencing for individual bins, for each of the six randomly isolated variants.

(C) Scatter plot shows the GFP intensities of yeast variants shared by the two biological replicates. Pearson’s correlation coefficient r and the corresponding *P* value are shown.

(D) The flowchart shows the number of variants passing individual computational cut-offs.

(E) The boxplot shows the GFP/dTomato ratio for variants with or without in-frame stop codons. *P* value was given by the Mann-Whitney *U* test.

(F) The boxplot shows the GFP/dTomato ratio for variants with or without uAUGs. *P* values were given by the Mann-Whitney *U* tests.

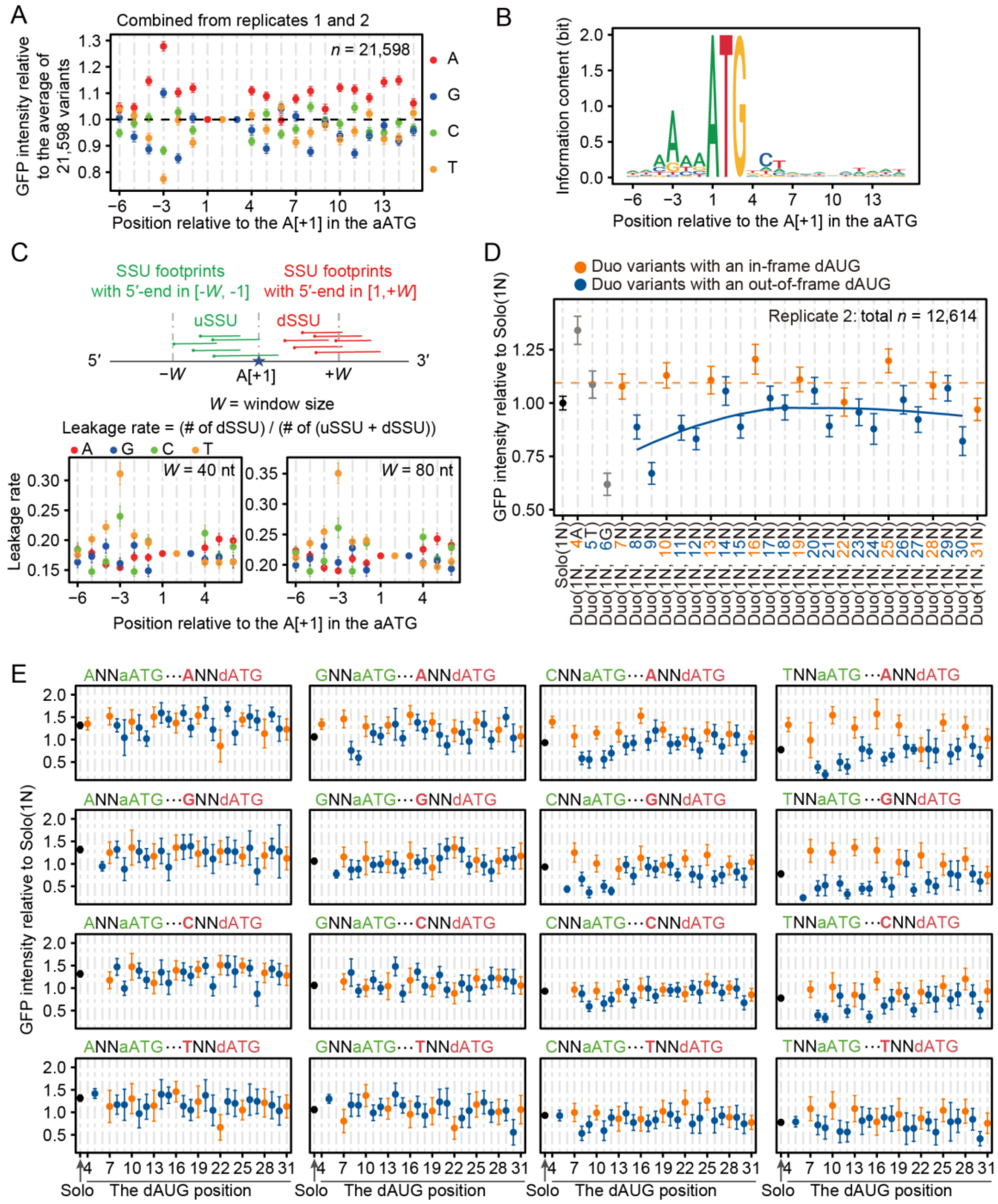

**Fig. S2. Frame- and context-dependent inhibitory effects on protein synthesis by proximal dAUGs, related to Fig. 1 and Fig. 2.**

(A) The average GFP intensity of yeast variants grouped by the nucleotide at each position. Variants were aligned according to the aATG. The GFP intensity for each variant was normalized by the average GFP intensity of all 21,598 variants. Error bars represent the 95% confidence intervals.

(B) Sequence logo around the aATG among the top 500 highly expressed genes in the yeast genome. The expression level data were retrieved from Nagalakshmi *et al*. (2008), and the sequence logo was generated with R package "ggseqlogo".

(C) The leakage rate estimated from the small ribosomal subunits (SSU) footprints on the yeast endogenous genes that were sequenced in Archer et al. (2016), according to the numbers of the SSU footprints with their 5′-end falling into the upstream window (uSSU) or the downstream window (dSSU) of the aATG. The results of two window sizes (W) are shown.

(D) The average GFP intensities (dots) and the 95% confidence intervals (error bars) of Duo variants measured in biological replicate 2. Similar to Fig. 1E.

(E) The average GFP intensities (dots) and the 95% confidence intervals (error bars) of Duo variants, grouped by the nucleotides at the −3 position of both aAUG and dAUG. Variants from both biological replicates were combined.

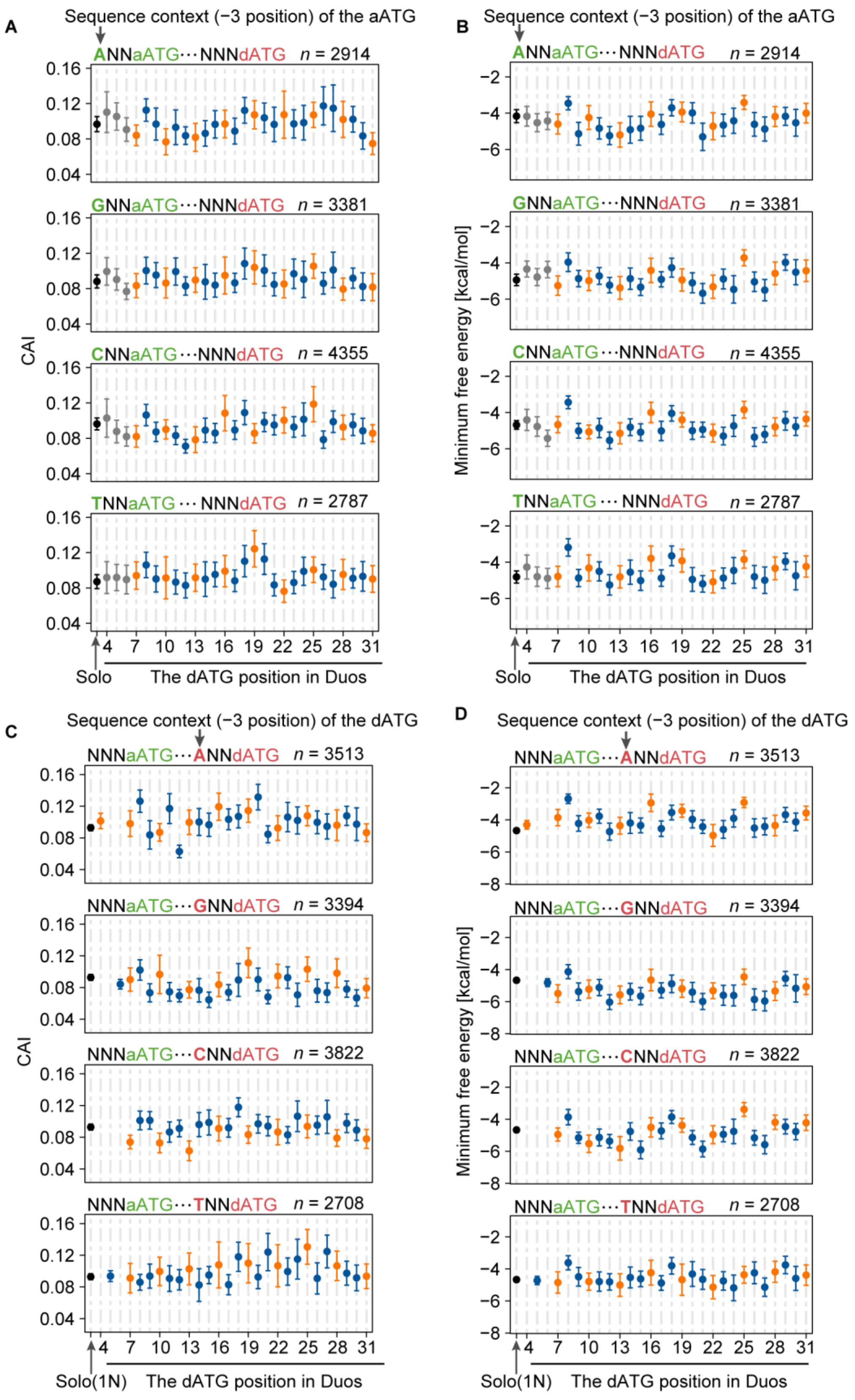


**Fig. S3. Codon adaption index (CAI) and minimum free energy (MFE) are not significantly varied among Duo variants with dATGs at various locations, related to Fig.2.**

Duo variants were grouped by the nucleotide at the −3 position of the aATG (A–B) or dATG (C–D). Error bars represent the 95% confidence intervals, and the total number of variants used in each panel is shown on the top. A higher CAI value means the tendency of using more preferred codons while a higher MFE value means more unstable mRNA secondary structure.

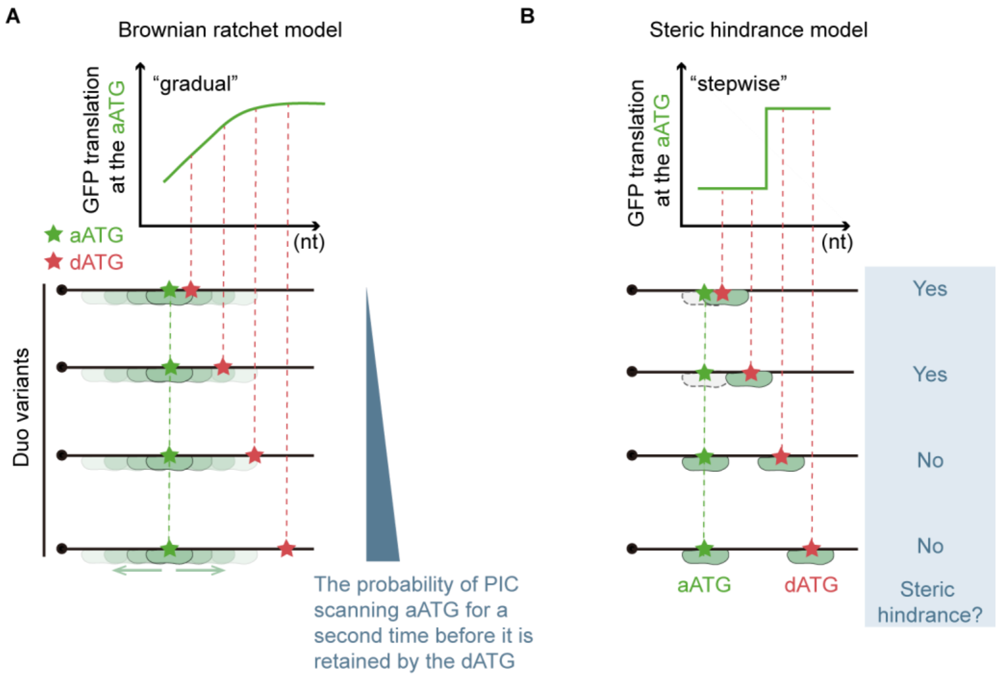


**Fig. S4. Two mechanistic explanations of the inhibitory effects on protein synthesis by proximal dAUGs, related to Fig. 2.**

(A) In the Brownian ratchet model, a proximal dAUG can retain PICs that miss the aAUG, reducing the chance for a second (or more) inspection of the aAUG in a distance-dependent manner. This model predicts that the inhibitory effects of dAUG will gradually decrease with the distance increase between aATG and dATG.

(B) In the steric hindrance model, the PIC occupying at a nearby dAUG (e.g., waiting for the ribosomal subunit joining) could hinder the aAUG recognition by the PIC (illustrated by the dashed lines). This model predicts that the inhibitory effects of a dAUG should depend on particular patterns of spacing between aATG and dATG, resulting in a stepwise transition in response to that distance.

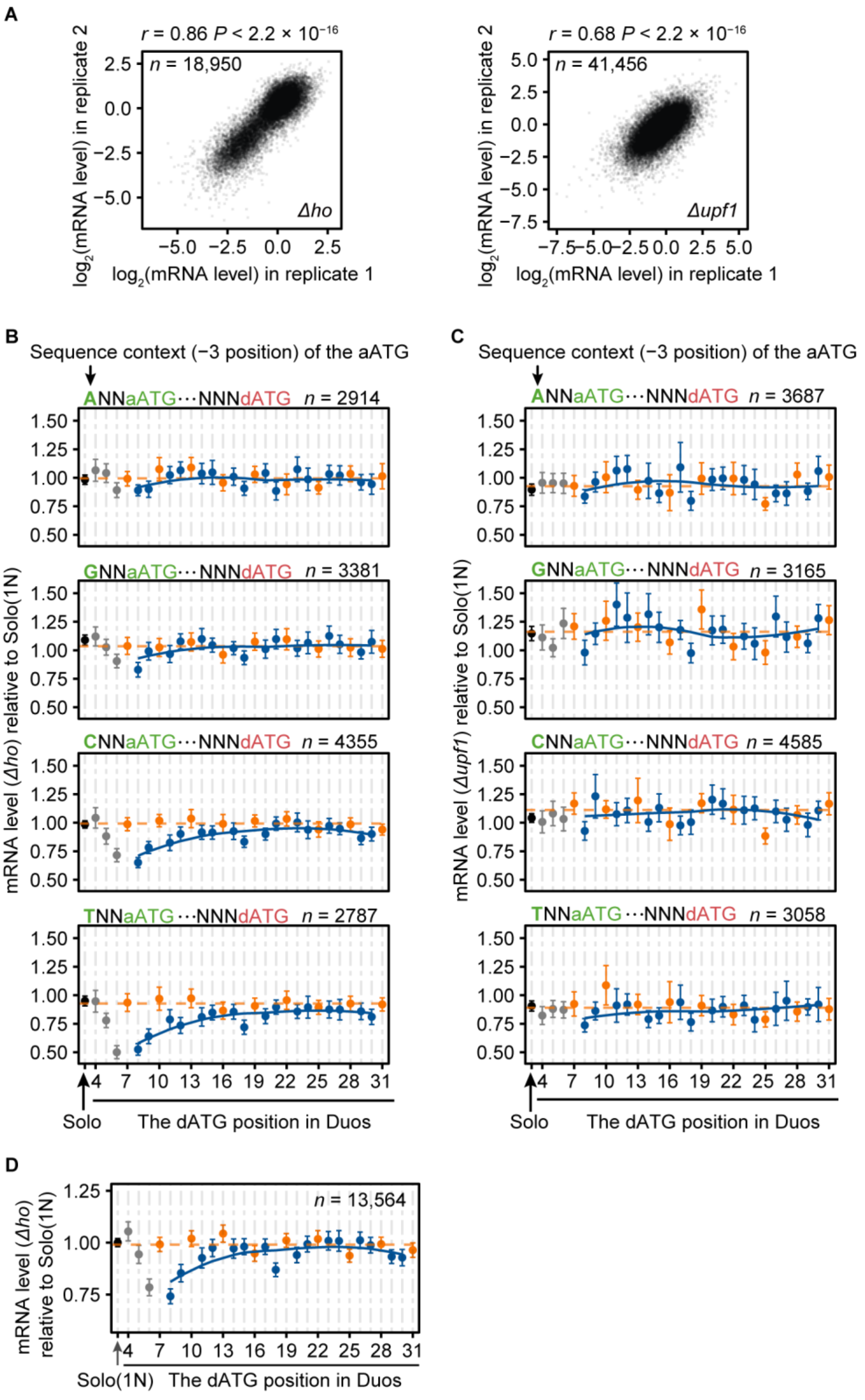


**Fig. S5. High-throughput measurement of mRNA levels for dATG variants, related to Fig. 3.**

(A) Scatter plots showing mRNA levels of dATG variants that were measured in both biological replicates, in the genetic background of *Δho* or *Δupf1*.

(B–C) The average mRNA levels (dots) and the 95% confidence intervals (error bars) of Duo variants, in the background of *Δho* (B) and *Δupf1* (C), grouped according to sequence contexts of the aATG.

(D) The average mRNA levels (dots) and the 95% confidence intervals (error bars) of Duo variants that do not contain out-of-frame stop codons corresponding to dATGs in the variable region.

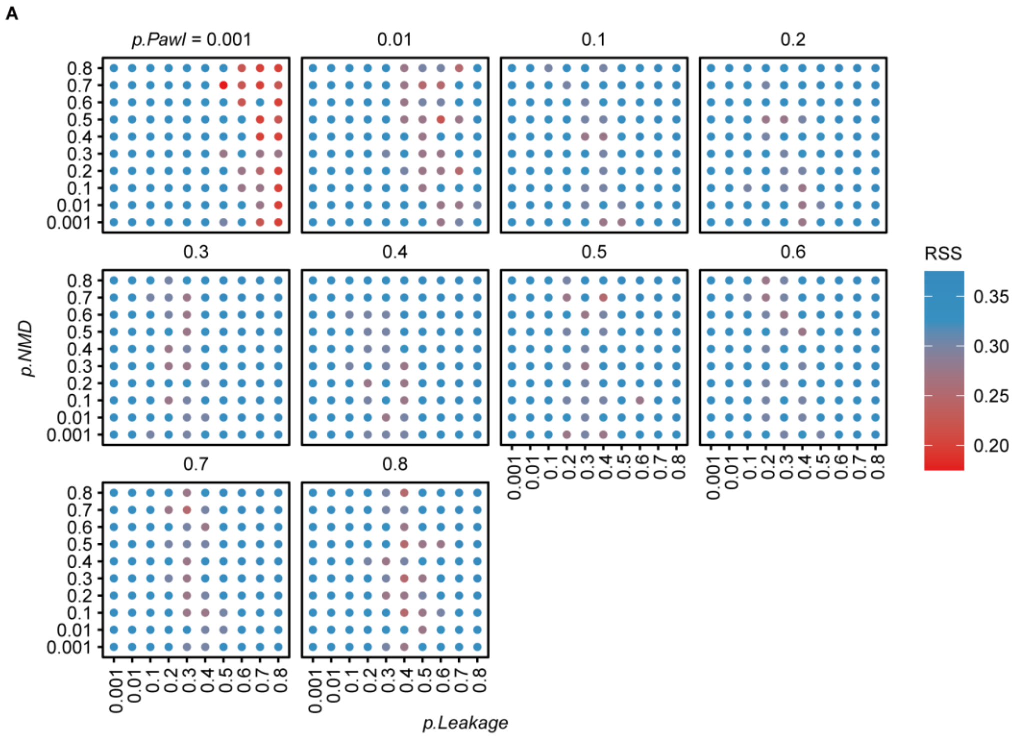


B

| Parameter | Median | Mean | Standard deviation |
|---|---|---|---|
| *p.Pawl* | 0.001 | 0.001 | 0.000 |
| *p.Leakage* | 0.75 | 0.73 | 0.09 |
| *p.NMD* | 0.55 | 0.51 | 0.26 |

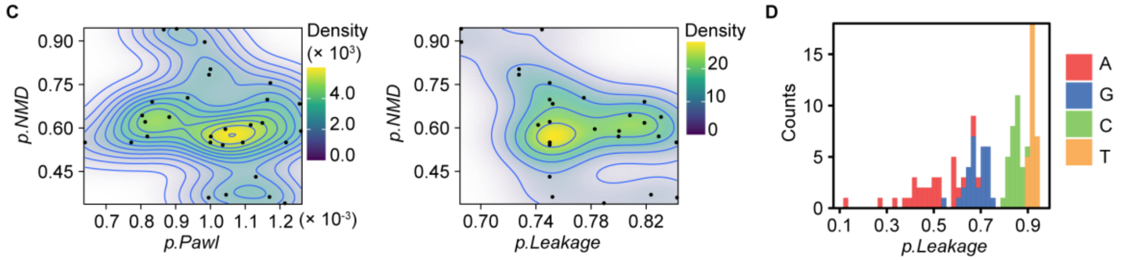


**Fig. S6. The optimization of the parameters in the Brownian ratchet scanning model using the MCMC algorithms, related to Fig. 5.**

(A) The residual sum of squares (RSS, values shown in color) estimated from the observed GFP intensities in the yeast experiments and the simulated protein expression level, using each of the 1000 parameter sets (*p.Pawl*, *p.Leakage*, and *p.NMD*).

(B) Table shows the summary statistics for *p.Pawl*, *p.Leakage*, and *p.NMD*, for the top 10 (out of 1000) parameter sets according to the RSS.

(C) Similar to Fig. 5E, two two-dimensional density plots show additional distributions of the outcome parameter values among the 30 MCMC chains.

(D) Stacked histogram plot shows the distributions of the outcome values of *p.Leakage* for ATGs in the A-, G-, C-, and T-contexts, respectively, among 30 MCMC chains.

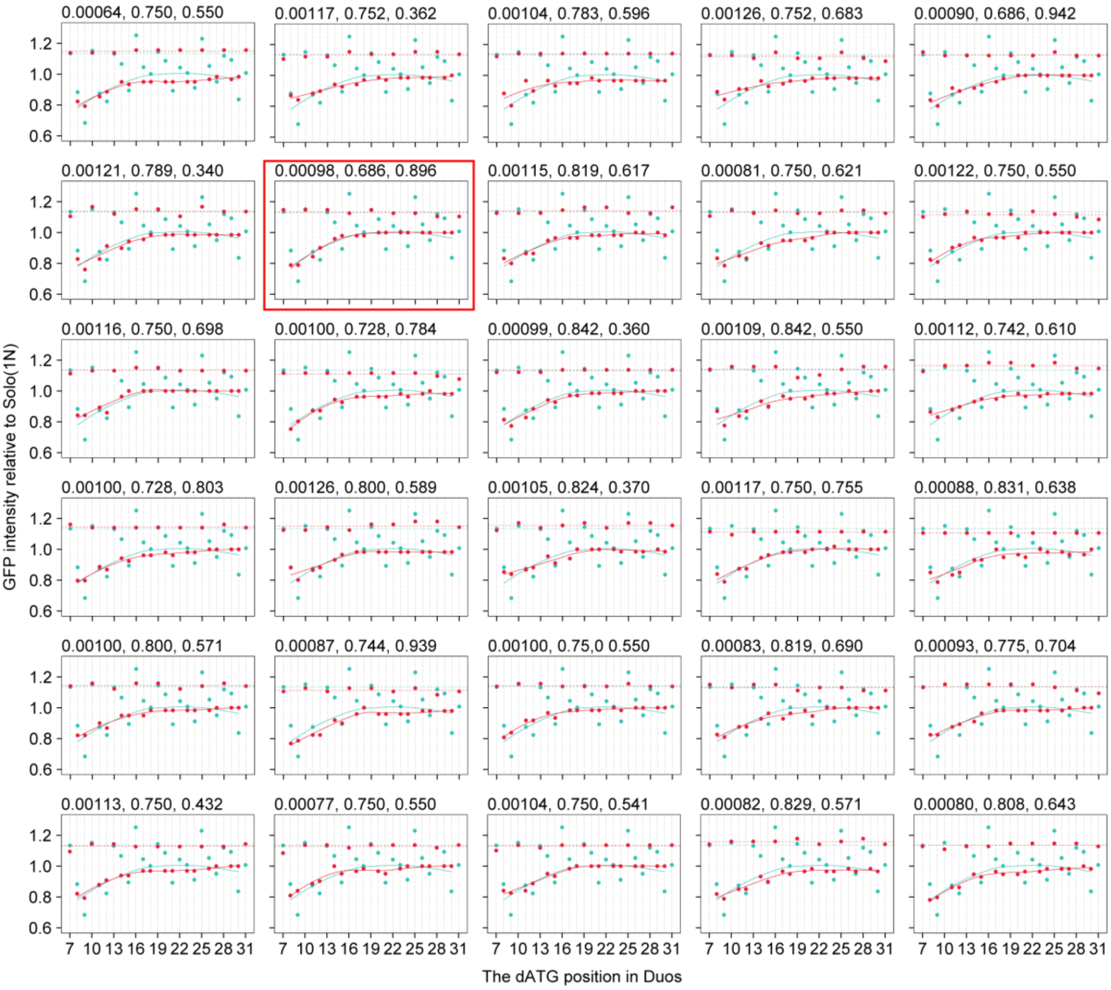


**Fig. S7. The comparison of the observed GFP intensities of Duo variants in the yeast experiments and the simulated GFP intensities for each of the 30 sets of optimized parameters by the MCMC algorithms, related to Fig. 5.**

Similar to Fig. 5D, showing the results of all 30 MCMC chains. The optimized values of parameters are shown on the top of each panel, in the order of *p.Pawl*, *p.Leakage*, and *p.NMD*. The result presented in Fig. 5D is shown in the red box.

### Supplemental Tables

**Table S1. Doped nucleotide oligos used to construct the yeast dATG library, related to Fig. 1, Fig. S1,and STAR Methods.**

| Oligo name | Sequence (5′-3′)[a] | Fraction used to mix oligos |
|---|---|---|
| 6NATG30N_random R1 | TAACCCGGGGATCCGTCGACCTGCNNNNNNNNNNNNNNNNNNNNNNNNNNNCATCATNNNNNNTTTTTCTCCTTGACGTTAAAGTA | 0.03 |
| 6NATG30N_random R2 | TAACCCGGGGATCCGTCGACCTGCNNNNNNNNNNNNNNNNNNNNNNNNNNCATNCATNNNNNNTTTTTCTCCTTGACGTTAAAGTA | 0.03 |
| 6NATG30N_random R3 | TAACCCGGGGATCCGTCGACCTGCNNNNNNNNNNNNNNNNNNNNNNNNNCATNNCATNNNNNNTTTTTCTCCTTGACGTTAAAGTA | 0.03 |
| 6NATG30N_random R4 | TAACCCGGGGATCCGTCGACCTGCNNNNNNNNNNNNNNNNNNNNNNNNCATNNNCATNNNNNNTTTTTCTCCTTGACGTTAAAGTA | 0.03 |
| 6NATG30N_random R5 | TAACCCGGGGATCCGTCGACCTGCNNNNNNNNNNNNNNNNNNNNNNNCATNNNNCATNNNNNNTTTTTCTCCTTGACGTTAAAGTA | 0.03 |
| 6NATG30N_random R6 | TAACCCGGGGATCCGTCGACCTGCNNNNNNNNNNNNNNNNNNNNNNCATNNNNNCATNNNNNNTTTTTCTCCTTGACGTTAAAGTA | 0.03 |
| 6NATG30N_random R7 | TAACCCGGGGATCCGTCGACCTGCNNNNNNNNNNNNNNNNNNNNNCATNNNNNNCATNNNNNNTTTTTCTCCTTGACGTTAAAGTA | 0.03 |
| 6NATG30N_random R8 | TAACCCGGGGATCCGTCGACCTGCNNNNNNNNNNNNNNNNNNNNCATNNNNNNNCATNNNNNNTTTTTCTCCTTGACGTTAAAGTA | 0.03 |
| 6NATG30N_random R9 | TAACCCGGGGATCCGTCGACCTGCNNNNNNNNNNNNNNNNNNNCATNNNNNNNNCATNNNNNNTTTTTCTCCTTGACGTTAAAGTA | 0.03 |
| 6NATG30N_random R10 | TAACCCGGGGATCCGTCGACCTGCNNNNNNNNNNNNNNNNNNCATNNNNNNNNNCATNNNNNNTTTTTCTCCTTGACGTTAAAGTA | 0.03 |
| 6NATG30N_random R11 | TAACCCGGGGATCCGTCGACCTGCNNNNNNNNNNNNNNNNNCATNNNNNNNNNNCATNNNNNNTTTTTCTCCTTGACGTTAAAGTA | 0.03 |
| 6NATG30N_random R12 | TAACCCGGGGATCCGTCGACCTGCNNNNNNNNNNNNNNNNCATNNNNNNNNNNNCATNNNNNNTTTTTCTCCTTGACGTTAAAGTA | 0.03 |

| Oligo name | Sequence (5′-3′)[a] | Fraction used to mix oligos |
|---|---|---|
| 6NATG30N_random R13 | TAACCCGGGGATCCGTCGACCTGCNNNNNNNNNNNNNNN<br>CATNNNNNNNNNNNNCATNNNNNNNTTTTTCTCCTTGACGT<br>TAAAGTA | 0.03 |
| 6NATG30N_random R14 | TAACCCGGGGATCCGTCGACCTGCNNNNNNNNNNNNNNC<br>ATNNNNNNNNNNNNNCATNNNNNNNTTTTTCTCCTTGACG<br>TTAAAGTA | 0.03 |
| 6NATG30N_random R15 | TAACCCGGGGATCCGTCGACCTGCNNNNNNNNNNNNNCA<br>TNNNNNNNNNNNNNNCATNNNNNNNTTTTTCTCCTTGACG<br>TTAAAGTA | 0.03 |
| 6NATG30N_random R16 | TAACCCGGGGATCCGTCGACCTGCNNNNNNNNNNNNCAT<br>NNNNNNNNNNNNNNNCATNNNNNNNTTTTTCTCCTTGACG<br>TTAAAGTA | 0.03 |
| 6NATG30N_random R17 | TAACCCGGGGATCCGTCGACCTGCNNNNNNNNNNNCATN<br>NNNNNNNNNNNNNNNCATNNNNNNNTTTTTCTCCTTGACG<br>TTAAAGTA | 0.03 |
| 6NATG30N_random R18 | TAACCCGGGGATCCGTCGACCTGCNNNNNNNNNNCATNN<br>NNNNNNNNNNNNNNNCATNNNNNNNTTTTTCTCCTTGACG<br>TTAAAGTA | 0.03 |
| 6NATG30N_random R19 | TAACCCGGGGATCCGTCGACCTGCNNNNNNNNNCATNNN<br>NNNNNNNNNNNNNNNCATNNNNNNNTTTTTCTCCTTGACG<br>TTAAAGTA | 0.03 |
| 6NATG30N_random R20 | TAACCCGGGGATCCGTCGACCTGCNNNNNNNNCATNNNN<br>NNNNNNNNNNNNNNNCATNNNNNNNTTTTTCTCCTTGACG<br>TTAAAGTA | 0.03 |
| 6NATG30N_random R21 | TAACCCGGGGATCCGTCGACCTGCNNNNNNNCATNNNNN<br>NNNNNNNNNNNNNNNCATNNNNNNNTTTTTCTCCTTGACG<br>TTAAAGTA | 0.03 |
| 6NATG30N_random R22 | TAACCCGGGGATCCGTCGACCTGCNNNNNNCATNNNNNN<br>NNNNNNNNNNNNNNNCATNNNNNNNTTTTTCTCCTTGACG<br>TTAAAGTA | 0.03 |
| 6NATG30N_random R23 | TAACCCGGGGATCCGTCGACCTGCNNNNNCATNNNNNNN<br>NNNNNNNNNNNNNNNCATNNNNNNNTTTTTCTCCTTGACG<br>TTAAAGTA | 0.03 |
| 6NATG30N_random R24 | TAACCCGGGGATCCGTCGACCTGCNNNNCATNNNNNNNN<br>NNNNNNNNNNNNNNNCATNNNNNNNTTTTTCTCCTTGACG<br>TTAAAGTA | 0.03 |
| 6NATG30N_random R25 | TAACCCGGGGATCCGTCGACCTGCNNNCATNNNNNNNNN<br>NNNNNNNNNNNNNNNCATNNNNNNNTTTTTCTCCTTGACG<br>TTAAAGTA | 0.03 |

| Oligo name | Sequence (5′-3′)[a] | Fraction used to mix oligos |
|---|---|---|
| 6NATG30N_random R26 | TAACCCGGGGATCCGTCGACCTGCNNCATNNNNNNNNNN<br>NNNNNNNNNNNNNNNCATNNNNNNTTTTTCTCCTTGACG<br>TTAAAGTA | 0.03 |
| 6NATG30N_random R27 | TAACCCGGGGATCCGTCGACCTGCNCATNNNNNNNNNNN<br>NNNNNNNNNNNNNNNCATNNNNNNTTTTTCTCCTTGACG<br>TTAAAGTA | 0.03 |
| 6NATG30N_random R28 | TAACCCGGGGATCCGTCGACCTGCCATNNNNNNNNNNNN<br>NNNNNNNNNNNNNNNCATNNNNNNTTTTTCTCCTTGACG<br>TTAAAGTA | 0.03 |
| 6NATG30N_random R29 | TAACCCGGGGATCCGTCGACCTGCNNNNNNNNNNNNNNN<br>NNNNNNNNNNNNNNNCATNNNNNNTTTTTCTCCTTGACG<br>TTAAAGTA | 0.15 |

**Table S2. Primers used in this study, related to STAR Methods.**

| Experiment | Usage | Primer name | Sequence (5′-3′) |
|---|---|---|---|
| **Construction of BY4742-dTomato yeast strain** | Construction | Gal7-del F | TATTATGCAGAGCATCAACATGATAAAAAAAAACAGTTGAATATTCCCTCAAAAATGGGTCGACGGATCCCCGGG |
| | | Gal7-del R | AAACCAGGCAGTTAATAGAAAAAATATGATATGAATGAATATTCCACTTTCTTTATCGATGAATTCGAGCTCG |
| | Verification | Gal7-colony F | TATTATGCAGAGCATCAACATGATA |
| | | Gal7-colony R | AAACCAGGCAGTTAATAGAAAAAAT |
| **Construction of the yeast dATG library** | Construction | Gal1 upstream F | ACATGGCATTACCACCATATACAT |
| | | GFP fusion F | GCAGGTCGACGGATCCCCGG |
| | | Gal1 downstream R | CCGGTCTAGGAAATCGGTGAAAGC |
| | Verification | Gal1 up colony F | CACTTTGTAACTGAGCTGTC |
| | | Gal1 down colony R | TCATTCTGATGTCATACGAC |
| **Construction of BY4742 *Δupf1* strain** | Construction | *Δupf1*::*natMX* F | AGGAAGGGCAGCAAGACCGAATATACTTTTTATATTACATCAATCATTGTCATTATCAAAGATCTGTTTAGCTTGCCTT |
| | | *Δupf1*::*natMX* R | TTGAGCCGTTTTGTATCACAAGCCAAGTTTAACATTTTATTTTAACAGGGTTCACCGAAATCGATGAATTCGAGCTCGT |
| | Verification | *Δupf1* colony F | TTGGGAGGGACACCTTTATAC |
| | | *Δupf1* colony R | TTGCAGTGCGCCGTAAAGAG |
| **Construction of BY4742 *Δho* strain** | Construction | *Δho*::*natMX* F | ACTATTAGCTCTAAATCCATATCCTCATAAGCAGCAATCAATTCTATCTATACTTTAAAAGATCTGTTTAGCTTGCCTT |
| | | *Δho*::*natMX* R | ATCCAAAATATTAAATTTTACTTTTATTACATACAACTTTTTAAACTAATATACACATTATCGATGAATTCGAGCTCGT |
| | Verification | *Δho* colony F | CAATTCCTATTCTAAATGGC |
| | | *Δho* colony R | TTTCTACTCCAGCATTCTAG |
| **Construction of Illumina sequencing library for FACS-seq** | The 1st round of PCR | Barcode501 | CACGACGCTCTTCCGATCTNNNNNNNNNNNNATCACGCCTCTATACTTTAACGTCAAGG |
| | | Barcode502 | CACGACGCTCTTCCGATCTNNNNNNNNNNNNCGATGTCCTCTATACTTTAACGTCAAGG |
| | | Barcode503 | CACGACGCTCTTCCGATCTNNNNNNNNNNNNTTAGGCCCTCTATACTTTAACGTCAAGG |
| | | Barcode504 | CACGACGCTCTTCCGATCTNNNNNNNNNNNNTGACCACCTCTATACTTTAACGTCAAGG |
| | | Barcode505 | CACGACGCTCTTCCGATCTNNNNNNNNNNNNACAGTGCCTCTATACTTTAACGTCAAGG |
| | | Barcode506 | CACGACGCTCTTCCGATCTNNNNNNNNNNNNGCCAATCCTCTATACTTTAACGTCAAGG |

| Experiment | Usage | Primer name | Sequence (5′-3′) |
|---|---|---|---|
| | | Barcode507 | CACGACGCTCTTCCGATCTNNNNNNNNNNNNCAGATCCCTCTATACTTTAACGTCAAGG |
| | | Barcode508 | CACGACGCTCTTCCGATCTNNNNNNNNNNNNACTTGACCTCTATACTTTAACGTCAAGG |
| | | Barcode701 | CAGACGTGTGCTCTTCCGATCTNNNNNNNNNNNNNCGTGATTGTTAATTAACCCGGGGAT |
| | | Barcode702 | CAGACGTGTGCTCTTCCGATCTNNNNNNNNNNNNNACATCGTGTTAATTAACCCGGGGAT |
| | | Barcode703 | CAGACGTGTGCTCTTCCGATCTNNNNNNNNNNNNNGCCTAATGTTAATTAACCCGGGGAT |
| | | Barcode704 | CAGACGTGTGCTCTTCCGATCTNNNNNNNNNNNNNTGGTCATGTTAATTAACCCGGGGAT |
| | | Barcode705 | CAGACGTGTGCTCTTCCGATCTNNNNNNNNNNNNNCACTGTTGTTAATTAACCCGGGGAT |
| | | Barcode706 | CAGACGTGTGCTCTTCCGATCTNNNNNNNNNNNNNATTGGCTGTTAATTAACCCGGGGAT |
| | | Barcode707 | CAGACGTGTGCTCTTCCGATCTNNNNNNNNNNNNNGATCTGTGTTAATTAACCCGGGGAT |
| | | Barcode708 | CAGACGTGTGCTCTTCCGATCTNNNNNNNNNNNNNTCAAGTTGTTAATTAACCCGGGGAT |
| | The 2nd round of PCR | P5 end-2 F | AATGATACGGCGACCACCGAGATCTACACTCTTTCCCTACACGACGCTCTTCCGATC |
| | | P7 end-2 index17 R (rep 1) | CAAGCAGAAGACGGCATACGAGATCTCTACGTGACTGGAGTTCAGACGTGTGCTCTTCC |
| | | P7 end-2 index18 R (rep 2) | CAAGCAGAAGACGGCATACGAGATGCGGACGTGACTGGAGTTCAGACGTGTGCTCTTCC |
| **Construction of Illumina sequencing library for RNA-seq** | The 1st round of PCR | Barcode501 (for *Δho* rep 1) | See the sequence of “Barcode501” |
| | | Barcode502 (for *Δho* rep 2 and *Δupf1* rep 2) | See the sequence of “Barcode502” |
| | | Barcode503 (for *Δupf1* rep 1) | See the sequence of “Barcode503” |
| | | Barcode701 (for *Δho* rep 1) | See the sequence of “Barcode701” |
| | | Barcode702 (for *Δho* rep 2 and *Δupf1* rep 2) | See the sequence of “Barcode702” |
| | | Barcode703 (for *Δupf1* rep 1) | See the sequence of “Barcode703” |
| | The 2nd round of PCR | P5 end-2 F | See the sequence of “P5 end-2 F” |
| | | P7 end-2 index39 R | CAAGCAGAAGACGGCATACGAGATGTATAGGTGACTGGAGTTCAGACGTGTGCTCTTCC |
| **Construction of Illumina** | The 1st round of PCR | Barcode501 (for *Δho* rep 1) | See the sequence of “Barcode501” |

| Experiment | Usage | Primer name | Sequence (5′-3′) |
|---|---|---|---|
| **sequencing library for DNA-seq** | | Barcode502 (for *Δho* rep 2 and *Δupf1* rep 2) | See the sequence of "Barcode502" |
| | | Barcode503 (for *Δupf1* rep 1) | See the sequence of "Barcode503" |
| | | Barcode701 (for *Δho* rep 1) | See the sequence of "Barcode701" |
| | | Barcode702 (for *Δho* rep 2 and *Δupf1* rep 2) | See the sequence of "Barcode702" |
| | | Barcode703 (for *Δupf1* rep 1) | See the sequence of "Barcode703" |
| | The 2nd round of PCR | P5 end-2 F | See the sequence of "P5 end-2 F" |
| | | P7 end-2 index38 R | CAAGCAGAAGACGGCATACGAGATAGCTA<br>GGTGACTGGAGTTCAGACGTGTGCTCTTCC |
| **Construction of overlapping dual-fluorescence reporter** | PCR of the fragment 1 | GFP-dTomato F | ACTATTAGCTCTAAATCCATATCCTCATAAG<br>CAGCAATCAATTCTATCTATACTTTAAATCA<br>TTATCAATACTCGCCAT |
| | | GFP-dTomato R-aaaATG | AAGCCCGGGGATCCGTCGACCTGCTTCATT<br>TTCCATAAAATTTCGAAACTAAGTTCTGGTG<br>T |
| | | GFP-dTomato R-tttATG | AAGCCCGGGGATCCGTCGACCTGCTTCATA<br>AACCATAAAATTTCGAAACTAAGTTCTGGT<br>GT |
| | PCR of the fragment 2 | GFP F | GCAGGTCGACGGATCCCCGG |
| | | GFP-dTomato R | ATCCAAAATATTAAATTTTACTTTTATTACA<br>TACAACTTTTTAAACTAATATACACATTCGG<br>GTAATAACTGATATAAT |
| | Fusion PCR | GFP-dTomato F | See the sequence of "GFP-dTomato F" |
| | | GFP-dTomato R | See the sequence of "GFP-dTomato R" |
| **Construction of dual-luciferase reporter** | Construction of the control variant | HeLa-accATG0 F | CAATCCGGTACTGTTGGTAAAAATTTTATG<br>GACCACGACCACGACCACGACGAAGATGCC<br>AAAAACATTAA |
| | | HeLa-accATG0 R | TTAATGTTTTTGGCATCTTCGTCGTGGTCGT<br>GGTCGTGGTCCATAAAATTTTTACCAACAG<br>TACCGGATTG |
| | Construction of the +8 dATG variant | HeLa-accATG8 F | CAATCCGGTACTGTTGGTAAAAATTTTATG<br>GACCATGACCACGACCACGACGAAGATGCC<br>AAAAACATTAA |
| | | HeLa-accATG8 R | TTAATGTTTTTGGCATCTTCGTCGTGGTCGT<br>GGTCATGGTCCATAAAATTTTTACCAACAG<br>TACCGGATTG |
| | Construction of the +14 dATG variant | HeLa-accATG14 F | CAATCCGGTACTGTTGGTAAAAATTTTATG<br>GACCACGACCATGACCACGACGAAGATGCC<br>AAAAACATTAA |

| Experiment | Usage | Primer name | Sequence (5′-3′) |
|---|---|---|---|
| | | HeLa-accATG14 R | TTAATGTTTTTGGCATCTTCGTCGTGGTCATGGTCGTGGTCCATAAAATTTTTACCAACAGTACCGGATTG |
| | Construction of the +20 dATG variant | HeLa-accATG20 F | CAATCCGGTACTGTTGGTAAAAATTTTATGGACCACGACCACGACCATGACGAAGATGCCAAAAACATTAA |
| | | HeLa-accATG20 R | TTAATGTTTTTGGCATCTTCGTCATGGTCGTGGTCGTGGTCCATAAAATTTTTACCAACAGTACCGGATTG |
| **Construction of yeast strains with dATG at different positions** | PCR of the fragment 1 | TEF F | ACTATTAGCTCTAAATCCATATCCTCATAAGCAGCAATCAATTCTATCTATACTTTAAACATAGCTTCAAAATGTTTCT |
| | | TEF R#8 | TAACCCGGGGATCCGTCGACCTGCTTCGTTTTCGTTTTCGTTTTCATTTTCCATAAAATTCTTAGATTAGATTGCTATGC |
| | | TEF R#14 | TAACCCGGGGATCCGTCGACCTGCTTCGTTTTCGTTTTCATTTTCGTTTTCCATAAAATTCTTAGATTAGATTGCTATGC |
| | | TEF R#20 | TAACCCGGGGATCCGTCGACCTGCTTCGTTTTCATTTTCGTTTTCGTTTTCCATAAAATTCTTAGATTAGATTGCTATGC |
| | | TEF R#26 | TAACCCGGGGATCCGTCGACCTGCTTCATTTTCGTTTTCGTTTTCGTTTTCCATAAAATTCTTAGATTAGATTGCTATGC |
| | PCR of the fragment 2 | GFP fusion-1F | GCAGGTCGACGGATCCCCGGGTTAATTAACAGTAAAGGAGAAGAACTTTTC |
| | | GFP R | TCCAAAATATTAAATTTTACTTTTATTACATACAACTTTTTAAACTAATATACACATTCGGGTAATAACTGATATAAT |
| | Fusion PCR | TEF F | See the sequence of “TEF F” |
| | | GFP R | See the sequence of “GFP R” |

**Table S3. Coding sequences for the dual-fluorescence reporter and the overlapping dual-fluorescence reporter, related to Figs. 1F, Figs. 2C, and STAR Methods.**

| Name | Sequence (5′-3′) |
|---|---|
| TEF promoter-*GFP* CDS-*CYC1* terminator-*TDH3* promoter-*dTomato* CDS-*ADH1* terminator-*URA3MX* [a] | CATAGCTTCAAAATGTTTCTACTCCTTTTTTACTCTTCCAGATTTTCTCGGACTCC<br>GCGCATCGCCGTACCACTTCAAAACACCCAAGCACAGCATACTAAATTTCCCCT<br>CTTTCTTCCTCTAGGGTGTCGTTAATTACCCGTACTAAAGGTTTGGAAAAGAAA<br>AAAGAGACCGCCTCGTTTCTTTTTCTTCGTCGAAAAAGGCAATAAAAATTTTTA<br>TCACGTTTCTTTTTCTTGAAAATTTTTTTTTTGATTTTTTTCTCTTTCGATGACCTC<br>CCATTGATATTTAAGTTAATAAACGGTCTTCAATTTCTCAAGTTTCAGTTTCATT<br>TTTCTTGTTCTATTACAACTTTTTTTACTTCTTGCTCATTAGAAAGAAAGCATAG<br>CAATCTAATCTAAGTTTATGAGTAAAGGAGAAGAACTTTTCACTGGAGTTGTCC<br>CAATTCTTGTTGAATTAGATGGTGATGTTAATGGGCACAAATTTTCTGTCAGTG<br>GAGAGGGTGAAGGTGATGCAACATACGGAAAACTTACCCTTAAATTTATTTGCA<br>CTACTGGAAAACTACCTGTTCCATGGCCAACACTTGTCACTACTTTCACTTATGG<br>TGTTCAATGCTTTTCAAGATACCCAGATCATATGAAACGGCATGACTTTTTCAA<br>GAGTGCCATGCCCGAAGGTTATGTACAGGAAAGAACTATATTTTTCAAAGATGA<br>CGGGAACTACAAGACACGTGCTGAAGTCAAGTTTGAAGGTGATACCCTTGTTAA<br>TAGAATCGAGTTAAAAGGTATTGATTTTAAAGAAGATGGAAACATTCTTGGACA<br>CAAATTGGAATACAACTATAACTCACACAATGTATACATCATGGCAGACAAAC<br>AAAAGAATGGAATCAAAGTTAACTTCAAAATTAGACACAACATTGAAGATGGA<br>AGCGTTCAACTAGCAGACCATTATCAACAAAATACTCCAATTGGCGATGGCCCT<br>GTCCTTTTACCAGACAACCATTACCTGTCCACACAATCTGCCCTTTCGAAAGATC<br>CCAACGAAAAGAGAGACCACATGGTCCTTCTTGAGTTTGTAACAGCTGCTGGGA<br>TTACACATGGCATGGATGAACTATACAAATAGTCATGTAATTAGTTATGTCACG<br>CTTACATTCACGCCCTCCCCCCACATCCGCTCTAACCGAAAAGGAAGGAGTTAG<br>ACAACCTGAAGTCTAGGTCCCTATTTATTTTTTTATAGTTATGTTAGTATTAAGA<br>ACGTTATTTATATTTCAAATTTTTCTTTTTTTTCTGTACAGACGCGTGTACGCATG<br>TAACATTATACTGAAAACCTTGCTTGAGAAGGTTTTGGGACGCTCGAAGTCATT<br>ATCAATACTCGCCATTTCAAAGAATACGTAAATAATTAATAGTAGTGATTTTCC<br>TAACTTTATTTAGTCAAAAAATTAGCCTTTTAATTCTGCTGTAACCCGTACATGC<br>CCAAAATAGGGGGCGGGTTACACAGAATATATAACATCGTAGGTGTCTGGGTG<br>AACAGTTTATTCCTGGCATCCACTAAATATAATGGAGCCCGCTTTTTAAGCTGG<br>CATCCAGAAAAAAAAAGAATCCCAGCACCAAAATATTGTTTTCTTCACCAACCA<br>TCAGTTCATAGGTCCATTCTCTTAGCGCAACTACAGAGAACAGGGGCACAAACA<br>GGCAAAAAACGGGCACAACCTCAATGGAGTGATGCAACCTGCCTGGAGTAAAT<br>GATGACACAAGGCAATTGACCCACGCATGTATCTATCTCATTTTCTTACACCTTC<br>TATTACCTTCTGCTCTCTCTGATTTGGAAAAAGCTGAAAAAAAAAGGTTGAAACC<br>AGTTCCCTGAAATTATTCCCCTACTTGACTAATAAGTATATAAAGACGGTAGGT<br>ATTGATTGTAATTCTGTAAATCTATTTCTTAAACTTCTTAAATTCTACTTTTATAG<br>TTAGTCTTTTTTTTAGTTTTAAAACACCAGAACTTAGTTTCGACGGATTCTAGAA<br>CTAGTGATGGTGAGCAAGGGCGAGGAGGTCATCAAAGAGTTCATGCGCTTCAA<br>GGTGCGCATGGAGGGCTCCATGAACGGCCACGAGTTCGAGATCGAGGGCGAGG<br>GCGAGGGCCGCCCCTACGAGGGCACCCAGACCGCCAAGCTGAAGGTGACCAAG |

| Name | Sequence (5′-3′) |
|---|---|
| | GGCGGCCCCCTGCCCTTCGCCTGGGACATCCTGTCCCCCCAGTTCATGTACGGC<br>TCCAAGGCGTACGTGAAGCACCCCGCCGACATCCCCGATTACAAGAAGCTGTCC<br>TTCCCCGAGGGCTTCAAGTGGGAGCGCGTGATGAACTTCGAGGACGGCGGTCT<br>GGTGACCGTGACCCAGGACTCCTCCCTGCAGGACGGCACGCTGATCTACAAGGT<br>GAAGATGCGCGGCACCAACTTCCCCCCCGACGGCCCCGTAATGCAGAAGAAGA<br>CCATGGGCTGGGAGGCCTCCACCGAGCGCCTGTACCCCCGCGACGGCGTGCTG<br>AAGGGCGAGATCCACCAGGCCCTGAAGCTGAAGGACGGCGGCCACTACCTGGT<br>GGAGTTCAAGACCATCTACATGGCCAAGAAGCCCGTGCAACTGCCCGGCTACT<br>ACTACGTGGACACCAAGCTGGACATCACCTCCCACAACGAGGACTACACCATC<br>GTGGAACAGTACGAGCGCTCCGAGGGCCGCCACCACCTGTTCCTGTACGGCATG<br>GACGAGCTGTACAAGTAAGGCGCGCCACTTCTAAATAAGCGAATTTCTTATGAT<br>TTATGATTTTTATTATTAAATAAGTTATAAAAAAAATAAGTGTATACAAATTTTA<br>AAGTGACTCTTAGGTTTTAAAACGAAAATTCTTATTCTTGAGTAACTCTTTCCTG<br>TAGGTCAGGTTGCTTTCTCAGGTATAGTATGAGGTCGCTCTTATTGACCACACCT<br>TCAATTCAATTCATCATTTTTTTTTTATTCTTTTTTTTGATTTCGGTTTCTTTGAAA<br>TTTTTTTGATTCGGTAATCTCCGAACAGAAGGAAGAACGAAGGAAGGAGCACA<br>GACTTAGATTGGTATATATACGCATATGTAGTGTTGAAGAAACATGAAATTGCC<br>CAGTATTCTTAACCCAACTGCACAGAACAAAAACCTGCAGGAAACGAAGATAA<br>ATCATGTCGAAAGCTACATATAAGGAACGTGCTGCTACTCATCCTAGTCCTGTT<br>GCTGCCAAGCTATTTAATATCATGCACGAAAAGCAAACAAACTTGTGTGCTTCA<br>TTGGATGTTCGTACCACCAAGGAATTACTGGAGTTAGTTGAAGCATTAGGTCCC<br>AAAATTTGTTTACTAAAAACACATGTGGATATCTTGACTGATTTTTCCATGGAG<br>GGCACAGTTAAGCCGCTAAAGGCATTATCCGCCAAGTACAATTTTTTACTCTTC<br>GAAGACAGAAAATTTGCTGACATTGGTAATACAGTCAAATTGCAGTACTCTGCG<br>GGTGTATACAGAATAGCAGAATGGGCAGACATTACGAATGCACACGGTGTGGT<br>GGGCCCAGGTATTGTTAGCGGTTTGAAGCAGGCGGCAGAAGAAGTAACAAAGG<br>AACCTAGAGGCCTTTTGATGTTAGCAGAATTGTCATGCAAGGGCTCCCTATCTA<br>CTGGAGAATATACTAAGGGTACTGTTGACATTGCGAAGAGCGACAAAGATTTT<br>GTTATCGGCTTTATTGCTCAAAGAGACATGGGTGGAAGAGATGAAGGTTACGAT<br>TGGTTGATTATGACACCCGGTGTGGGTTTAGATGACAAGGGAGACGCATTGGGT<br>CAACAGTATAGAACCGTGGATGATGTGGTCTCTACAGGATCTGACATTATTATT<br>GTTGGAAGAGGACTATTTGCAAAGGGAAGGGATGCTAAGGTAGAGGGTGAACG<br>TTACAGAAAAGCAGGCTGGGAAGCATATTTGAGAAGATGCGGCCAGCAAAACT<br>AAAAAACTGTATTATAAGTAAATGCATGTATACTAAACTCACAAATTAGAGCTT<br>CAATTTAATTATATCAGTTATTACCCTATGCGGTGTGAAATACCGCACAGATGC<br>GTAAGGAGAAAATACCGCATCAGG |

| Name | Sequence (5′-3′) |
| --- | --- |
| GFP-aaaATG-dTomato[b] | **(GFP)ATG**GAAA**(dTomato)ATG**AAGCAGGTCGACGGATCCCCGGGCTTATTAAC<br>AGTAAAGGAGAAGAACTTTTCACTGGAGTTGTCCCAATTCTTGTTGAATTGGAT<br>GGTGATGTTAATGGGCACAAATTTTCTGTCAGTGGAGAGGGTGAAGGTGATGC<br>AACATACGGAAAACTTACCCTTAAATTTATTTGCACTACTGGAAAACTACCTGT<br>TCCATGGCCAACACTTGTCACTACTTTCACTTATGGTGTTCAATGCTTTTCAAGA<br>TACCCAGATCATCTTAAACGGCATGACTTTTTCAAGAGTGCCATGCCCGAAGGT<br>TATGTACAGGAAAGAACTATATTTTTCAAAGATGACGGGAACTACAAGACACG<br>TGCTGAAGTCAAGTTTGAAGGTGATACCCTTGTTAATAGAATCGAGCTTAAAGG<br>TATTGATTTTAAAGAAGATGGAAACATTCTTGGACACAAATTGGAATACAACTA<br>TAACTCACACAATGTATACATCATGGCAGACAAACAAAAGAATGGAATCAAAG<br>TTAACTTCAAAATTAGACACAACATTGAAGATGGAAGCGTTCAATTGGCAGACC<br>ATTATCAACAAAATACTCCAATTGGCGATGGCCCTGTCCTTTTACCAGACAACC<br>ATTACCTGTCCACACAATCTGCCCTTTCGAAAGATCCCAACGAAAAGAGAGACC<br>ACATGGTCCTTCTTGAGTTTGTTACAGCTGCTGGGATTACACATGGCATGGATG<br>AACTATACAAATAGAggttctggtggtgctactaatttttctttgttgaaattggctggtgatgttgaattgaatccaggtcc<br>aATGGTGAGCAAGGGCGAGGAGGTCATCAAAGAGTTCATGCGCTTCAAGGTGC<br>GCATGGAGGGCTCCATGAACGGCCACGAGTTCGAGATCGAGGGCGAGGGCGAG<br>GGCCGCCCCTACGAGGGCACCCAGACCGCCAAGCTGAAGGTGACCAAGGGCGG<br>CCCCCTGCCCTTCGCCTGGGACATCCTGTCCCCCCAGTTCATGTACGGCTCCAAG<br>GCGTACGTGAAGCACCCCGCCGACATCCCCGATTACAAGAAGCTGTCCTTCCCC<br>GAGGGCTTCAAGTGGGAGCGCGTGATGAACTTCGAGGACGGCGGTCTGGTGAC<br>CGTGACCCAGGACTCCTCCCTGCAGGACGGCACGCTGATCTACAAGGTGAAGA<br>TGCGCGGCACCAACTTCCCCCCCGACGGCCCCGTAATGCAGAAGAAGACCATG<br>GGCTGGGAGGCCTCCACCGAGCGCCTGTACCCCCGCGACGGCGTGCTGAAGGG<br>CGAGATCCACCAGGCCCTGAAGCTGAAGGACGGCGGCCACTACCTGGTGGAGT<br>TCAAGACCATCTACATGGCCAAGAAGCCCGTGCAACTGCCCGGCTACTACTACG<br>TGGACACCAAGCTGGACATCACCTCCCACAACGAGGACTACACCATCGTGGAA<br>CAGTACGAGCGCTCCGAGGGCCGCCACCACCTGTTCCTGTACGGCATGGACGA<br>GCTGTACAAGTAA |
| GFP-tttATG-dTomato[c] | **(GFP)ATG**GTTT**(dTomato)ATG**AAGCAGGTCGACGGATCCCCGGGCTTATTAACA<br>GTAAAGGAGAAGAACTTTTCACTGGAGTTGTCCCAATTCTTGTTGAATTGGATG<br>GTGATGTTAATGGGCACAAATTTTCTGTCAGTGGAGAGGGTGAAGGTGATGCA<br>ACATACGGAAAACTTACCCTTAAATTTATTTGCACTACTGGAAAACTACCTGTT<br>CCATGGCCAACACTTGTCACTACTTTCACTTATGGTGTTCAATGCTTTTCAAGAT<br>ACCCAGATCATCTTAAACGGCATGACTTTTTCAAGAGTGCCATGCCCGAAGGTT<br>ATGTACAGGAAAGAACTATATTTTTCAAAGATGACGGGAACTACAAGACACGT<br>GCTGAAGTCAAGTTTGAAGGTGATACCCTTGTTAATAGAATCGAGCTTAAAGGT<br>ATTGATTTTAAAGAAGATGGAAACATTCTTGGACACAAATTGGAATACAACTAT<br>AACTCACACAATGTATACATCATGGCAGACAAACAAAAGAATGGAATCAAAGT<br>TAACTTCAAAATTAGACACAACATTGAAGATGGAAGCGTTCAATTGGCAGACC<br>ATTATCAACAAAATACTCCAATTGGCGATGGCCCTGTCCTTTTACCAGACAACC<br>ATTACCTGTCCACACAATCTGCCCTTTCGAAAGATCCCAACGAAAAGAGAGACC |

| Name | Sequence (5′-3′) |
| --- | --- |
| | ACATGGTCCTTCTTGAGTTTGTTACAGCTGCTGGGATTACACATGGCATGGATG<br>AACTATACAAATAGAggttctggtggtgctactaatttttctttgttgaaattggctggtgatgttgaattgaatccaggtcc<br>aATGGTGAGCAAGGGCGAGGAGGTCATCAAAGAGTTCATGCGCTTCAAGGTGC<br>GCATGGAGGGCTCCATGAACGGCCACGAGTTCGAGATCGAGGGCGAGGGCGAG<br>GGCCGCCCCTACGAGGGCACCCAGACCGCCAAGCTGAAGGTGACCAAGGGCGG<br>CCCCCTGCCCTTCGCCTGGGACATCCTGTCCCCCCAGTTCATGTACGGCTCCAAG<br>GCGTACGTGAAGCACCCCGCCGACATCCCCGATTACAAGAAGCTGTCCTTCCCC<br>GAGGGCTTCAAGTGGGAGCGCGTGATGAACTTCGAGGACGGCGGTCTGGTGAC<br>CGTGACCCAGGACTCCTCCCTGCAGGACGGCACGCTGATCTACAAGGTGAAGA<br>TGCGCGGCACCAACTTCCCCCCCGACGGCCCCGTAATGCAGAAGAAGACCATG<br>GGCTGGGAGGCCTCCACCGAGCGCCTGTACCCCCGCGACGGCGTGCTGAAGGG<br>CGAGATCCACCAGGCCCTGAAGCTGAAGGACGGCGGCCACTACCTGGTGGAGT<br>TCAAGACCATCTACATGGCCAAGAAGCCCGTGCAACTGCCCGGCTACTACTACG<br>TGGACACCAAGCTGGACATCACCTCCCACAACGAGGACTACACCATCGTGGAA<br>CAGTACGAGCGCTCCGAGGGCCGCCACCACCTGTTCCTGTACGGCATGGACGA<br>GCTGTACAAGTAA |

[a] The coding sequences of wild type *GFP*, *dTomato*, and *URA3* are shown in the 5′-3′ order (underlined).

[b, c] The translation initiation codons of GFP and dTomato are shown in bold, with the corresponding gene shown in parentheses. The modified *GFP* CDS and *dTomato* CDS are shown in the 5′-3′ order (underlined); the sequence of 2A self-cleaving peptide is shown in lowercase letters.

**Table S4. Number of variants in each sample, related to STAR Methods.**

| Strain | Biological replicate | Library | # of read pairs[a] | # of identified variants | Total number of Solo and Duo variants |
|---|---|---|---|---|---|
| *Δho* | replicate 1 | RNA-seq | 5,210,827 | 140,708 | 11,879 |
| | | DNA-seq | 2,641,781 | 119,716 | |
| | | FACS-seq | 7,978,902 | 170,698 | |
| | replicate 2 | RNA-seq | 5,022,016 | 134,709 | 12,614 |
| | | DNA-seq | 1,337,264 | 99,052 | |
| | | FACS-seq | 17,303,437 | 232,375 | |
| *Δupf1* | replicate 1 | RNA-seq | 5,696,430 | 108,147 | 16,225 |
| | | DNA-seq | 6,759,896 | 120,162 | |
| | replicate 2 | RNA-seq | 4,563,203 | 98,624 | 16,179 |
| | | DNA-seq | 6,225,186 | 115,747 | |

[a] Number of read pairs that passed the three criteria described in STAR Methods.

**Table S5. Information for individual bins in FACS-seq, related to Fig. 1, Fig. S1, and STAR Methods.**

| Biological replicate | Bin (*i*) | Number of cells collected in each bin | The median GFP/dTomato ($G_i$) | Proportion of cells belonging to the "gate" of each bin ($P_i$) |
|---|---|---|---|---|
| replicate 1 | bin 1 | 313,926 | 47.20 | 0.125 |
| | bin 2 | 181,218 | 29.91 | 0.070 |
| | bin 3 | 247,469 | 21.63 | 0.095 |
| | bin 4 | 394,686 | 14.19 | 0.099 |
| | bin 5 | 234,655 | 7.98 | 0.060 |
| | bin 6 | 156,297 | 3.42 | 0.040 |
| | bin 7 | 196,999 | 1.11 | 0.051 |
| | bin 8 | 350,000 | 0.25 | 0.460 |
| replicate 2 | bin 1 | 328,449 | 49.49 | 0.140 |
| | bin 2 | 168,373 | 29.61 | 0.071 |
| | bin 3 | 226,829 | 21.75 | 0.093 |
| | bin 4 | 350,000 | 14.34 | 0.088 |
| | bin 5 | 220,151 | 8.11 | 0.052 |
| | bin 6 | 151,088 | 3.60 | 0.039 |
| | bin 7 | 188,055 | 1.12 | 0.046 |
| | bin 8 | 350,000 | 0.24 | 0.471 |
